# Cryo-EM structures of Na^+^-pumping NADH-ubiquinone oxidoreductase from *Vibrio cholerae*

**DOI:** 10.1101/2022.04.19.488743

**Authors:** Jun-ichi Kishikawa, Moe Ishikawa, Takahiro Masuya, Masatoshi Murai, Yuki Kitazumi, Nicole L. Butler, Takayuki Kato, Blanca Barquera, Hideto Miyoshi

## Abstract

The Na^+^-pumping NADH-ubiquinone oxidoreductase (Na^+^-NQR) couples electron transfer from NADH to ubiquinone with Na^+^-pumping, generating an electrochemical Na^+^ gradient that is essential for energy-consuming reactions in bacteria. Since Na^+^-NQR is exclusively found in prokaryotes, it is a promising target for highly selective antibiotics. However, the molecular mechanism of inhibition is not well-understood for lack of the atomic structural information about an inhibitor-bound state. Here we present cryo-electron microscopy structures of Na^+^- NQR from *Vibrio cholerae* with or without a bound inhibitor at 2.5- to 3.1-Å resolution. The structures reveal the arrangement of all six redox cofactors including riboflavin, whose position has been under debate, and a newly assigned 2Fe-2S^NqrD/E^ cluster located between the membrane embedded NqrD and NqrE subunits. A large part of the hydrophilic NqrF near the cytoplasmic membrane surface is barely visible in the density map, suggesting a high degree of flexibility. This flexibility may be responsible to reducing the long distance between the 2Fe- 2S centers in NqrF and NqrD/E, consistent with physiologically relevant electron transfer. Two different types of specific inhibitors (korormicin A and aurachin D-42) bind to the N-terminal region of NqrB, which is disordered in the absence of inhibitors. The current inhibitor-bound structures reasonably explain our previous biochemical findings obtained by different chemistry-based experiments. This study provides a definite foundation for understanding the function of Na^+^-NQR and the molecular mechanism of its specific inhibitors to support molecular design of new antibiotics targeting the enzyme.

## INTRODUCTION

The Na^+^-pumping NADH-ubiquinone (UQ) oxidoreductase (Na^+^-NQR) couples electron transfer from NADH to UQ with Na^+^-pumping, generating an electrochemical Na^+^ gradient across the inner bacterial membrane, which is critical for energy-consuming reactions such as rotation of the flagellar motor, ion homeostasis, and uptake of nutrients (*1–3*). Na^+^-NQR is the first enzyme in the respiratory chain of many pathogenic bacteria such as *Vibrio cholerae*, *Vibrio alginolyticus*, and *Haemophilus influenzae*. This enzyme is an intrinsic membrane protein complex composed of six subunits (NqrA–F), encoded by the *nqr* operon, with a total molecular mass about 200 kDa, and contains at least five redox cofactors (FAD, a 2Fe-2S cluster, two covalently bound FMNs, and riboflavin). There is a consensus that the electron transfer takes place through the enzyme following the pathway: NADH → FAD^NqrF^ → 2Fe-2S^NqrF^ → FMN^NqrC^ → FMN^NqrB^ → riboflavin^NqrB^ → UQ (*4–13*). Since Na^+^-NQR is exclusively found in prokaryotes and is structurally unrelated to mitochondrial H^+^-pumping NADH-UQ oxidoreductase (respiratory complex I), it is a promising target for highly selective antibiotics (*3, 14, 15*).

Steuber et al. (*16*) reported an X-ray crystallographic structure of selenomethionine-labeled *V*. *cholerae* Na^+^-NQR at 3.5-Å resolution, which was built by placing the crystallographic structures of individual soluble domains of NqrA, NqrC, and NqrF into the low-resolution experimental electron density map. This structure provided valuable information about the overall architecture of the enzyme, though it included no bound UQ or inhibitor. The distances between several pairs of redox cofactors in the crystallographic structure (e.g. between FMN and riboflavin of NqrB) are too long (29–32 Å (edge-to-edge)) to support physiologically relevant electron transfer (*16, 17*). Therefore, it has been suggested that the subunits harboring the cofactors undergo large conformational changes to decrease these spatial gaps during catalytic turnover (*16*). In fact, the linker regions between the single transmembrane helix, the N-terminal ferredoxin-like domain, and the FAD-binding domain in NqrF were barely visible in the electron density map, suggesting a high degree of flexibility. In addition, the crystallographic study modeled an extra cofactor, a single iron coordinated by four cysteine residues, two in NqrD (Cys29 and Cys112) and two in NqrE (Cys28 and Cys120), that was proposed to be an extra redox carrier, facilitating electron transfer between 2Fe-2S^NqrF^ to FMN^NqrC^. However, the presence of this iron center and its redox function remain to be verified by further experiments.

Based on photoaffinity labeling experiments (*18, 19*), we previously showed that korormicin A (a highly specific inhibitor of Na^+^-NQR, *18*, *20*) and aurachin D-type inhibitors (naphthoquinone-like compounds) bind to a part of the protruding N-terminal stretch starting with transmembrane helix (TMH) 1 of NqrB (Try23−Lys54, Supplementary Figure 1) and that the UQ head-ring binds to the cytoplasmic region of NqrA close to the N-terminal stretch of NqrB (Leu32−Met39 and Phe131−Lys138, Supplementary Figure 1). Thus, the N-terminal stretch of NqrB (Met1−Lys54) may be critical for regulating the UQ reaction at the adjacent NqrA. However, since approximately three-quarters (Met1−Pro37) of the protruding N- terminal stretch of NqrB was not modeled in the crystallographic structure (*16*), the structural and functional roles of this key segment remain unclear. Therefore, high-resolution structures of Na^+^-NQR are essential for understanding how all the cofactors are arranged in six subunits, how they participate in redox-coupled Na^+^ translocation, and how inhibitor molecules bind to the N-terminal region of NqrB.

Here, we present high-resolution (2.5–3.1 Å) structures of *V. cholerae* Na^+^-NQR in three different conditions determined by single-particle cryo-electron microscopy (EM): the native enzyme, without bound UQ or inhibitor, and enzymes with korormicin A or aurachin D-42 bound to NqrB. Although the structure of the N-terminal region of NqrB is disordered in the absence of inhibitor, its structure with bound korormicin A or aurachin D-42 could be fully assigned, including the missing N-terminal stretch of NqrB, because the binding of the inhibitors made the region rigid. As we previously predicted, based on chemistry-based studies (*21*), the N-terminal region orients itself toward the membrane phase by folding up at the cytoplasmic membrane surface rather than protruding into cytoplasmic medium along the surface of NqrA. Furthermore, our structures reveal that FMN^NqrC^, FMN^NqrB^, and riboflavin^NqrB^ embedded in the membrane phase, are closer to one another than in the X-ray crystallographic structure (*16*). The present study provides new insights not only into the overall structural rearrangements of Na^+^-NQR but also into the molecular mechanism of specific inhibitors.

## RESULTS

### Overall cryo-EM structure of Na^+^-NQR without bound inhibitor

The recombinant Na^+^-NQR from *V. cholerae* was expressed in a *V. cholerae* strain lacking the genomic *nqr* operon and purified by affinity chromatography in the presence of *n*-dodecyl- β-D-maltoside (DDM). Using the purified Na^+^-NQR, we determined its cryo-EM density map at 2.7-Å overall resolution (Extended Data Figures 1 and 2). The initial map was of sufficient quality to build an atomic model of most part of the enzyme. However, the density map of the region in the cytoplasmic NqrF subunit, facing the membrane-embedded subunits (NqrE and NqrD), was relatively weak and blurry compared to other parts of the enzyme. This was also the case for the X-ray crystallographic structure (*16*). We can confidently attribute this weak density to structural flexibility of this region of NqrF, rather than loss of the subunit from the complex, because we have confirmed that our preparation of Na^+^-NQR is intact and active, as judged by NADH-UQ_1_ oxidoreductase activity that is fully sensitive to inhibitor, and the presence of all six subunits (NqrA to NqrF) on SDS-PAGE (Extended Data Figure 3).

To obtain the structure of the cytoplasmic domain of NqrF, focused 3D classification without angular alignment, following local refinement on the NqrA and NqrB subunits was carried out. As expected, the cytoplasmic domain of NqrF showed high flexibility compared to other subunits (Supplementary Video 1). The focus 3D classification resulted in 10 classes; we selected the three, in which NqrF was clearly observed (Extended Data Figure 1). The particles belonging to the three classes were applied further Non-uniform refinement. As a result, we obtained high resolution structures of states 1, 2, and 3 from 82,790, 80,882, and 72,234 particles, respectively, at 3.1-Å resolution (Extended Data Figure 1). Atomic models of the cytoplasmic domain of NqrF of the three states were built from the resulting density maps. Figure 1A and 1B show the density map and the atomic model of Na^+^-NQR, respectively, taking state 1 as a representative. The atomic models for three states are almost identical except for the flexible cytoplasmic domain of NqrF. Details of data processing and statistics for the maps and models are summarized in Extended Data Figure 1 and Supplementary Tables 1 and 2.

**Figure 1:**
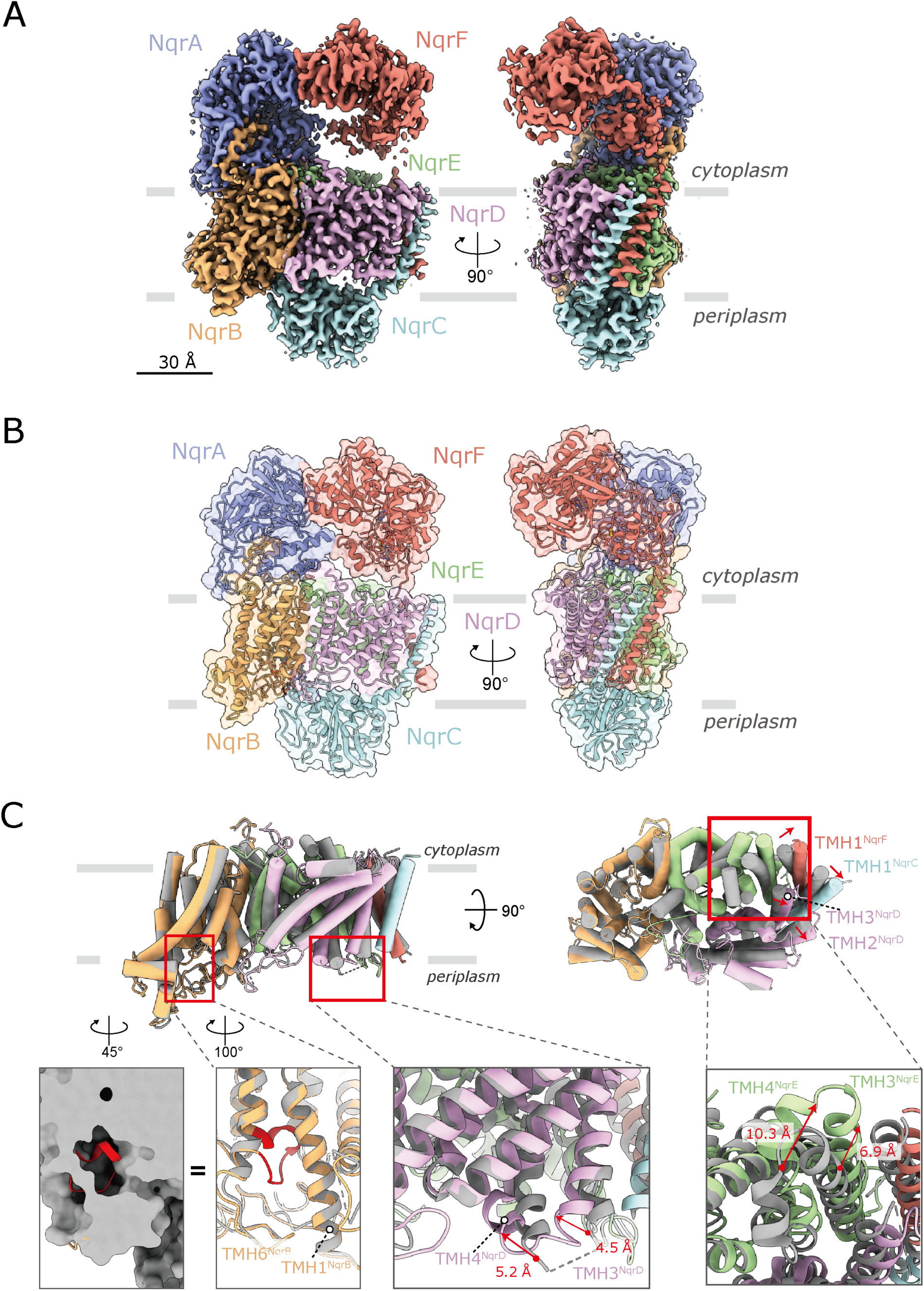
Overall structure of *V. cholerae* Na^+^-NQR. The density map (**A**) and the atomic model (**B**) of Na^+^-NQR, showing the six subunits NqrA–F. The density map and the atomic model are represented as semi-transparent surface and cartoon, respectively. The hydrophilic NqrA (*blue*) contains no TMH. NqrC (*pale*) and NqrF (*red*) are hydrophilic proteins, each anchored by a single TMH. The NqrB (*yellow*), NqrD (*pink*), and NqrE (*green*) subunits are integral membrane proteins. The membrane plane is indicated by gray lines. (**C**) Overlay of cryo-EM structure of the membrane domain (*color*) with the crystallographic structure (*gray*, *16*). NqrB, NqrC, NqrD, NqrE, and NqrF subunits have 10, 1, 6, 6, and 1 TMHs, respectively. TMHs are shown as *cylinder* models. The membrane domain contains a total of 24 TMHs. In the leftmost inset, the membrane interior loop (Gly266–Ser276) in the cryo-EM structure is indicated in *red*. The hydrophilic domain of each subunit, which protrudes from the membrane, has been deleted for clarity. TMHs are numbered.

The final high-quality density map allowed us to build an essentially complete model of all six subunits containing 1894 residues (98% of the total), except for the disordered N-terminal region (Gly2−Leu26) of NqrB. However, this region was well modeled in the inhibitor-bound enzyme, as described later. We note that the N-terminal amino acid of mature NqrB was determined to be glycine, not methionine, by direct Edman degradation (*21*). The NqrB, NqrD and NqrE subunits are integral membrane proteins that have 10, 6, and 6 TMHs, respectively. NqrC and NqrF are hydrophilic proteins each anchored by single TMH. A hydrophilic NqrA contains no TMH. The present study clearly defined all 24 TMHs (Figure 1C). The structures of the membrane domain in our model are almost identical to those in the crystallographic structure (*16*). Nevertheless, there are some differences in the local structures of TMHs between the two models (Figure 1C). TMH 2^NqrD^, TMH 3^NqrD^, TMH 1^NqrC^, and TMH 1^NqrF^, which form a helix bundle, are slightly shifted together toward the outside of the enzyme compared to the crystallographic structure. TMH 3^NqrD^ and TMH 4^NqrD^ slightly bent toward the center of the enzyme. The cytosolic region of TMH 3^NqrE^ and TMH 4^NqrE^ is shifted about 10 Å toward the outside of the enzyme. Furthermore, a membrane interior loop (Gly266–Ser276) lies in a narrow central cavity of NqrB (*red* in the leftmost inset); although this region was modeled as a part of TMH 6 and its periplasmic loop in the crystallographic structure.

The crystallographic study could not resolve some parts of the loops connecting TMHs 1– 2, 5–6, 7–8 of NqrB, TMHs 2–3, 3–4, 4–5, 5–6 of NqrD, and TMHs 2–3 of NqrE. The C-terminal loop region of NqrA (Arg330–Val375), which is critical to attach this subunit to the membrane by anchoring to the cytoplasmic surface of NqrB, was also not modeled. The present study successfully resolves all of the loops. On the whole, the overall cryo-EM structure of Na^+^- NQR closely resembles the crystallographic structure (*16*). In fact, the *r.m.s.d.* values for each subunit between the two structures are less than 2.5 Å (Supplementary Table S3). Nevertheless, the most notable difference is that in our models, the periplasmic hydrophilic domain of NqrC, which accommodates FMN^NqrC^, significantly shifts towards NqrB compared to that in the crystallographic structure, as discussed later. Altogether, the present high-resolution cryo-EM structures resolves some critical unsettled questions in the crystallographic structure.

### Structural basis of anchoring the cytoplasmic hydrophilic subunits NqrA and NqrF to the membrane

A large region of the hydrophilic NqrF subunit facing the membrane-embedded subunits is barely visible in the density map, suggesting a high degree of flexibility. As described in the above section, focused 3D classification visualized the cytoplasmic domain of NqrF in three different classes. NqrF binds tightly to the neighboring hydrophilic NqrA via electrostatic interactions at the interface of these subunits, where it is complementarily charged, as shown in Figure 2 using the state 1 model. The positive patch in NqrA, formed by Arg40, Lys61, Lys62, Arg84, and R85, interacts with the negative patch in NqrF, formed by Glu368, Glu371, Glu372, Glu397, and Glu399. In addition, negative tips of NqrA (Glu443 and Glu445) are in close proximity to positive tips of NqrF (Lys100 and Arg104). These electrostatic interactions in the contact area are consistent across the three different classes of NqrF (Extended Data Figure 4), suggesting that this contact area serves as the anchor that allows flexible structural changes of the rest of the hydrophilic parts of the subunit. In the structure of NqrF we can identify the NADH binding site, FAD and the 2Fe-2S center. The edge-to-edge distance between FAD^NqrF^ and 2Fe-2S^NqrF^ is 10.8 Å, which is almost identical to that determined by the crystallographic structure (9.8 Å, *16*).

**Figure 2:**
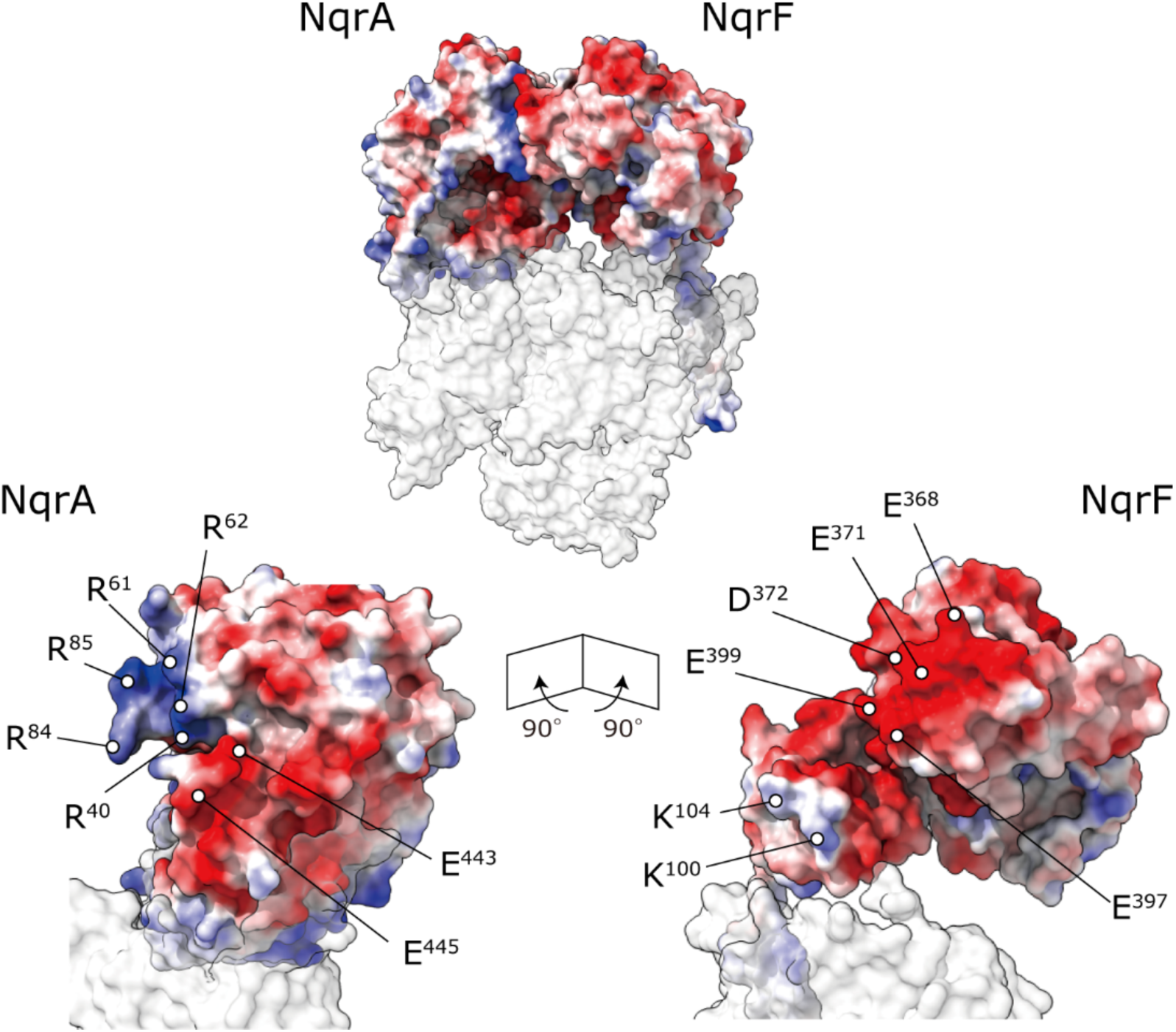
The cytoplasmic contact area between NqrA and NqrF. NqrF binds tightly to the neighboring hydrophilic NqrA via electrostatic interactions in the state 1 model. Lower panels show cross-section images of the contact area between the two subunits. The positively and negatively charged residues in the NqrA and NqrF are indicated in *blue* and *red*, respectively.

The hydrophilic NqrA subunit is attached to the cytoplasmic surface of the Na^+^-NQR complex through NqrB, but the details of this interaction have remained unclear because some parts of the interface between NqrA and NqrB were not resolved in the crystallographic structure (*16*). Elucidation of this issue is critical to understand the mechanism by which UQ accepts electrons from the enzyme, since the binding site of the UQ head-ring lies in NqrA (*8, 18*). Our cryo-EM structure reveals that the C-terminal loop region of NqrA (Arg330−Asn379) is intertwined with the almost all of the protruding part of the N-terminus of NqrB (Phe34−Leu54) at the cytoplasmic membrane surface (Extended Data Figure 5). In addition, a hydrophobic segment of the C-terminal loop of NqrA, that contains Leu334, Phe335, Try337, Ala338, Pro340, Leu352, and Leu355, would be inserted into the membrane phase *in vivo*. Together, these interactions anchor NqrA to the membrane surface.

### Revisiting the assignment of the redox cofactors

The intrinsic membrane subunits NqrD and NqrE are homologous proteins with a sequence identity of 37%. The X-ray crystallographic study reported that these two subunits form a two-fold pseudo-symmetrical dimer, resulting in the coordination of four highly conserved cysteine residues (NqrD-Cys29/NqrE-Cys120 and NqrE-Cys28/NqrD-Cys112) to a single iron atom at the center of the dimer surface. Although this iron atom appears to be critical for the electron transfer mediating between 2Fe-2S^NqrF^ and FMN^NqrC^, the presence of this putative iron center has yet to be definitely verified by biochemical and biophysical characterizations (*13*). However, in our cryo-EM structure, the density map at the interface between NqrD and NqrE shows a rhombus-shape resembling that of a 2Fe-2S cluster (Figure 3), as observed in structures of other redox enzymes (*22, 23*). The density is coordinated by the two pairs of conserved cysteine residues described above, which are known as the typical ligands for a 2Fe-2S cluster (*22, 23*). These features strongly suggest that the density belongs to a 2Fe-2S cluster rather than a single iron atom; therefore, we have assigned it as 2Fe-2S^NqrD/E^. Based on site-directed mutagenesis, Fadeeva et al. (*24*) reported that these conserved cysteine residues are necessary for the correct folding and/or the stability of Na^+^-NQR from *Vibrio harvevi*. The edge-to-edge distances between 2Fe-2S ^NqrD/E^ and FMN^NqrC^ in our model and between Fe^NqrD/E^ and FMN^NqrC^ in the crystallographic structure (*16*) are 29.1 and 7.9 Å, respectively. Since the positions of 2Fe-2S ^NqrD/E^ and Fe^NqrD/E^ in the membrane phase are almost identical between the two structural models, this remarkable difference in the distance is due to shift of the hydrophilic domain of NqrC, in which FMN^NqrC^ is bound, as described later. As the distances between 2Fe-2S ^NqrD/E^ and FMN^NqrC^ is too long to support physiologically relevant electron transfer, the subunits harboring the two cofactors would undergo a large conformational changes to reduce the spatial gap during the catalytic turnover.

**Figure 3:**
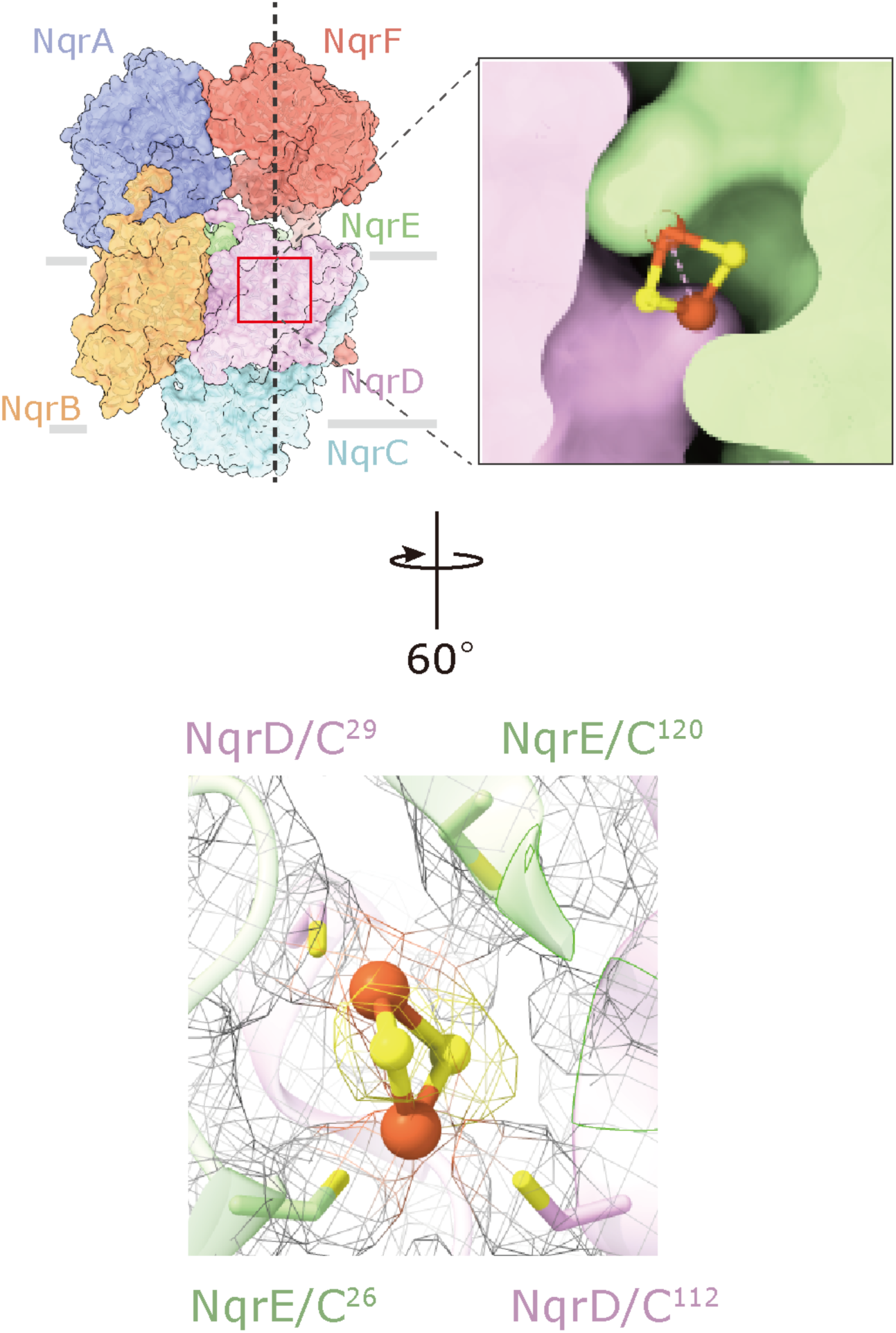
A new 2Fe-2S center located between subunits NqrD and NqrE. The density map at the interface between NqrD and NqrE resembles that of a 2Fe-2S cluster rather than a single iron atom. (Upper panel) The 2Fe-2S cluster at the center of the two-fold symmetrical axis of NqrD and NqrE. (Lower panel) The two pairs of conserved cysteine residues (NqrD-Cys29/NqrE-Cys120 and NqrE-Cys28 and NqrD-Cys112) coordinate the 2Fe-2S cluster. The Cys residues and 2Fe-2S cluster are shown as sticks and ball-and-sticks models, respectively. In this panel, the consensus map for Na^+^-NQR is used and shown as a mesh representation.

Riboflavin, a unique cofactor of Na^+^-NQR that forms a stable neutral semiquinone radical in the as-isolated state of the enzyme (*6, 25*), mediates the final electron transfer from FMN^NqrB^ to UQ. Based on a biochemical study using the Na^+^-NQRs that lack individual subunits, Casutt et al. (*26*) reported that riboflavin is localized to subunit NqrB. On the other hand, the crystallographic study assigned the riboflavin to a large patch of *Fo-Fc* density between NqrB and NqrE because no additional density was observed inside NqrB (*16*). Thus, the position of riboflavin is still under debate. We can now definitely assign riboflavin to the density map inside NqrB, which is surrounded by central TMHs 1, 3, 5 and 8 (Figure 4A and 4B). The riboflavin is accommodated in NqrB with the isoalloxazine ring oriented towards the inside of the protein via multiple hydrogen bonds with conserved polar amino acid residues. Thr162^TMH3^ and Asn203^TMH5^ form hydrogen bonds with the O2 and O4/N3 atoms of the isoalloxazine ring, respectively (Figure 4C). The 3’-hydroxy group of the ribityl side chain is in the vicinity of the O2 and N1 atoms, suggesting this hydroxy group works as an intramolecular hydrogen bond donor to these atoms. The rest of the hydroxy groups in the ribityl chain (at the 2’-, 4’-, and 5’- positions) are close enough to Asn200^TMH5^ and Asp346^TMH8^ to form hydrogen bonds (Figure 4C).

**Figure 4:**
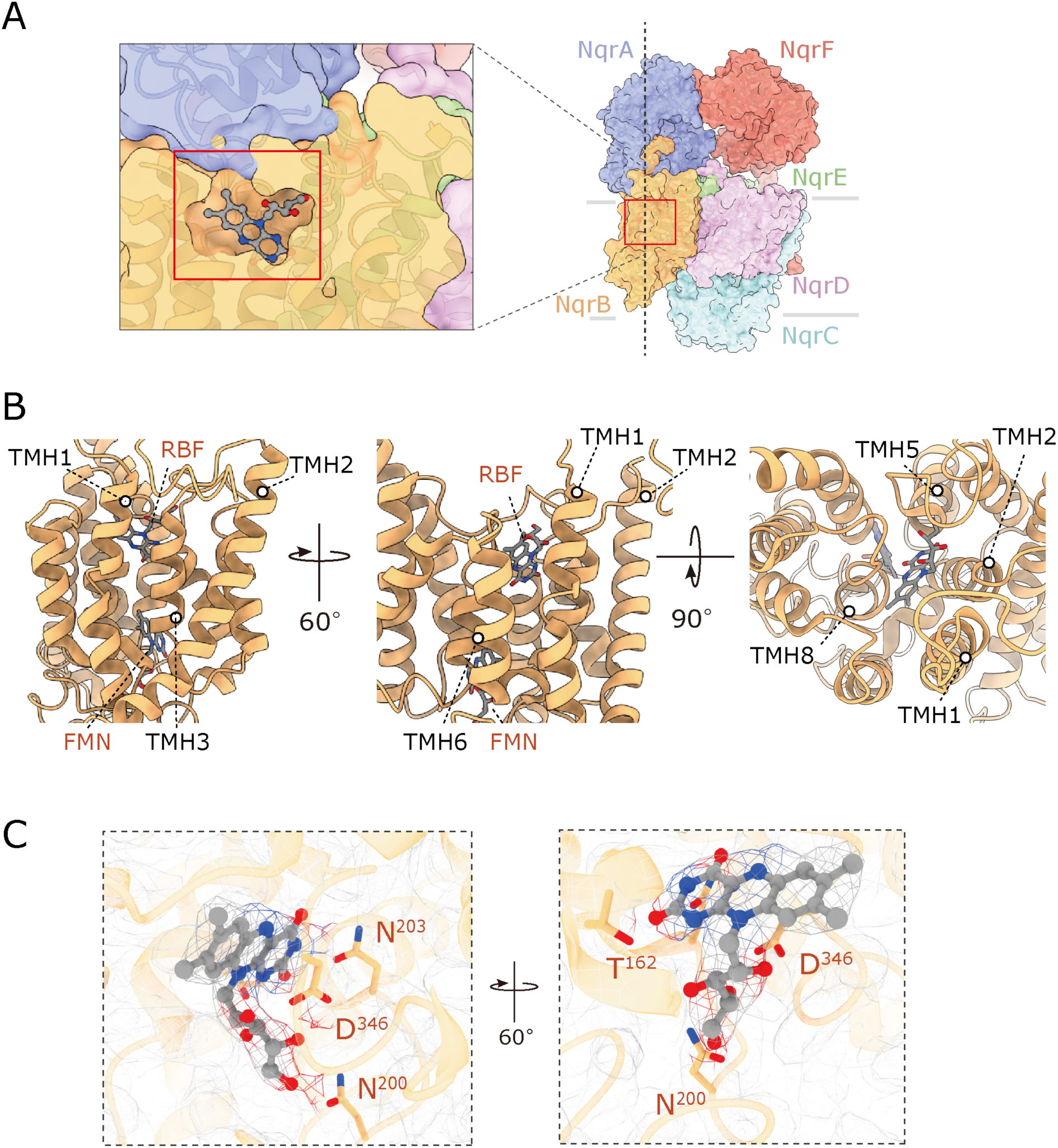
Riboflavin is located in NqrB. (**A**, **B**) The positions of riboflavin and FMN in NqrB are indicated. Riboflavin (*sticks* model) is surrounded by central TMHs 1, 3, 5, and 8 (*cartoon* model). (**C**) Close-up view of the binding site of riboflavin (*ball-and-sticks* model). The density map is shown as a *mesh* representation. The residues suggested to be involved in hydrogen bonds with riboflavin are indicated.

### Comparison of the positions of all cofactors between the cryo-EM and crystallographic structures

The NqrC subunit is a periplasmic protein anchored by a single TMH that is in close contact with the TMH 3 of NqrD and TMH 1 of NqrF (Figure 1C). The FMN^NqrC^ that accepts electrons from the 2Fe-2S^NqrD/E^ is covalently bound via a phosphoester bond to NqrC-Thr225 located in a helix of the C-terminus (Figure 5A). The body of FMN protrudes out from the NqrC protein matrix and lies in cavity formed by NqrB, NqrD, and NqrE on the periplasmic side of membrane. The isoalloxazine ring of the FMN is sandwiched between NqrC-Leu145 and -Leu176, and its N5 atom is likely to be stabilized by NqrC-Trp146 and -Thr173. Compared with the crystallographic structure (*16*), FMN^NqrC^ and these residues are substantially shifted toward FMN^NqrB^ and form a closed pocket at the interface between NqrB and NqrC, in which FMN^NqrC^ and FMN^NqrB^ face each other (Figure 5A). This arrangement reduces the edge-to-edge distance between the two cofactors to 7.8 Å, which is short enough to achieve efficient electron transfer. However, this same structural shift is also responsible for the long distance (29.1 Å) between 2Fe-2S^NqrD/E^ and FMN^NqrC^ in our model.

**Figure 5:**
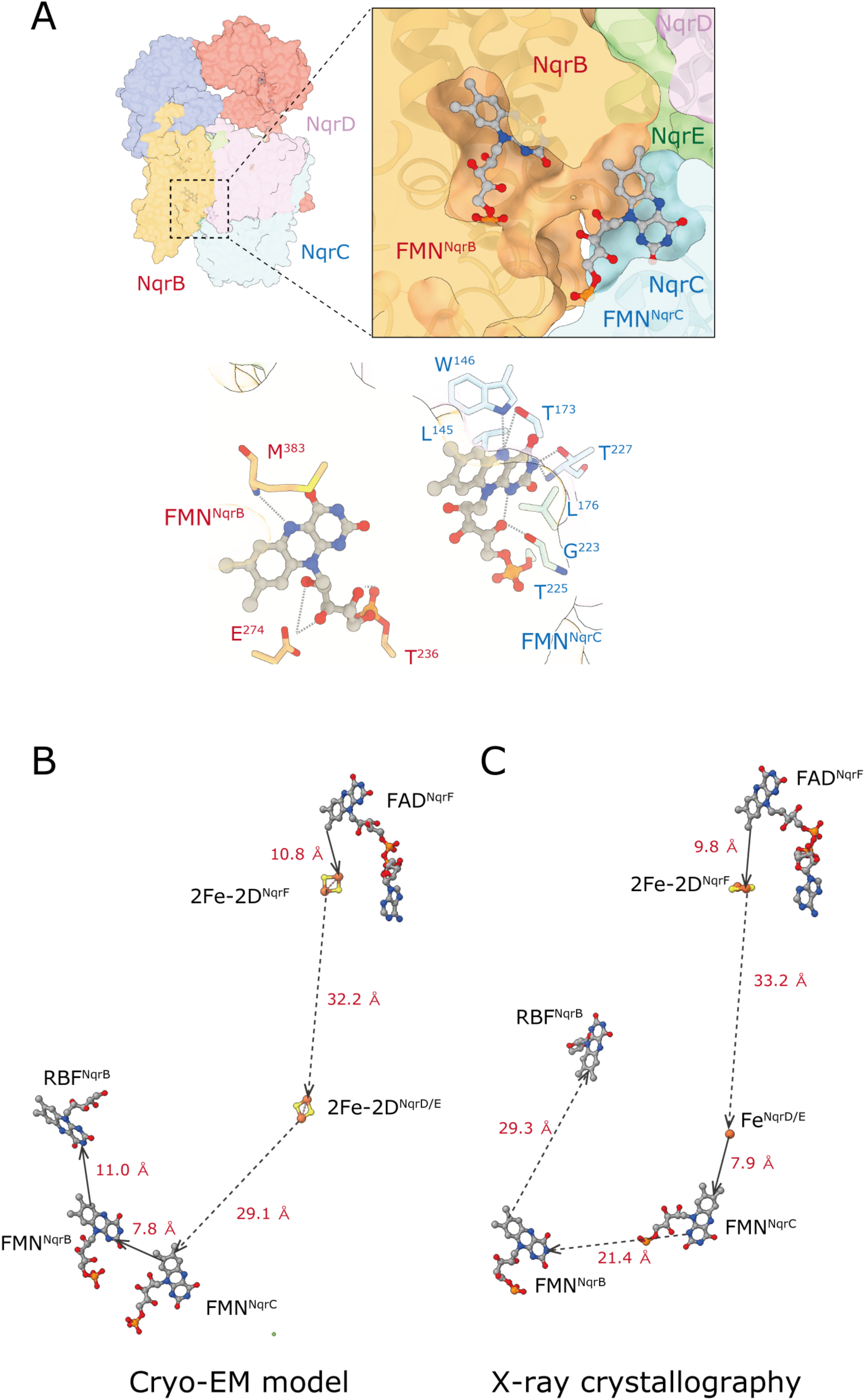
Positions of all cofactors in Na^+^-NQR. (**A**) The structure around FMN in NqrC. The head-group of FMN protrudes from the NqrC protein matrix and lies in the periplasmic medium cavity formed by NqrB, NqrD, and NqrE (*upper panel*). The residues that interact with FMN are indicated. FMN^NqrC^ and FMN^NqrB^ face each other in the closed pocket at the periplasmic interface between NqrB and NqrC (*lower panel*). The positions of all cofactors in the present cryo-EM model (**B**) and the X-ray crystallographic structure (**C**) (*16*) are indicated. The edge-to-edge distances of the cofactors are indicated.

The positions of all redox cofactors are compared between the present cryo-EM and X-ray crystallographic (*16*) structures in Figure 5B and 5C, although we have assigned the density between NqrD and NqrE as an 2Fe-2S cluster in place of a single iron. Even if we disregard the positions of the two cofactors (FAD and 2Fe-2S cluster) located in the very flexible NqrF for the time being, there is a pair of redox cofactors in each structure whose spatial distance is too long for efficient electron transfer, namely, 2Fe-2S^NqrD/E^ and FMN^NqrC^ and FMN^NqrB^ and riboflavin^NqrB^ in the cryo-EM and crystallographic structures, respectively. Nevertheless, as the position of riboflavin^NqrB^ was corrected in this study, the spatial distances between the three flavin cofactors (FMN^NqrC^, FMN^NqrB^, and riboflavin^NqrB^) are significantly shortened, consistent with efficient electron transfer between them. Clearly the catalytic cycle must involve significant structural changes that modulate the distances between cofactors, and are likely to be important for the coupling between redox and Na^+^ translocating reactions. Having structures of the fully-reduced and various partially-reduced states of Na^+^-NQR would be expected to be of great help in elucidating these important questions.

### Structures of Na^+^-NQR with bound inhibitor

To characterize the binding sites of inhibitors in *V. cholerae* Na^+^-NQR, we previously carried out photoaffinity labeling studies using korormicin and aurachin D-type derivatives (Supplementary Figure 1) (*18, 19*). The studies demonstrated that these inhibitors bind to the region Trp23−Lys54 in the N-terminus starting with TMH 1 of NqrB and/or a part of the cytoplasmic loop (NqrB-His153−Gly156) connecting TMHs 2*–*3 of NqrB. However, since most of the N-terminus of NqrB (Gly2−Pro37) could not be resolved in the crystallographic structure (*16*), leaving the molecular mechanism of the inhibitors unclear.

Therefore, we prepared cryo-EM grids of Na^+^-NQR in the presence of aurachin D-42 or korormicin A. Image processing provided the consensus density maps at 2.5- and 2.6-Å resolution for the aurachin D-42- and korormicin A-bound enzymes, respectively (Extended Data Figure 1B and 1C). However, the density corresponding to the hydrophilic domain of NqrF in these maps was faint, as was in the map for the enzyme without inhibitor (Extended Data Figures 1A and 2). Further 3D classification allowed us to obtain the final model at 3.0- and 2.9-Å resolution for the aurachin D-42- and korormicin A-bound enzymes, respectively (Supplementary Table 2). In both cases we successfully modeled the entire N-terminus of NqrB, in which the inhibitors are accommodated (Figure 6A and 6B). The loop connecting TMHs 2*–* 3 of NqrB forms a part of the rear wall of the binding cavity. These results are consistent with the results of our photoaffinity labeling studies (*18, 19*). The N-terminal region (Gly2−Leu26) of NqrB is flexible and disordered in the absence of inhibitor, as mentioned above, whereas the binding of inhibitor gives rise to the distinct conformation that we could identify by cryo-EM. It is noteworthy that the N-terminal region starting with TMH 1 first protrudes from the membrane phase and then turns back toward the membrane (bending at Arg43) and forms the binding cavity for inhibitors by folding up at the cytoplasmic membrane surface (Extended Data Figure 6A and 6B). Based on pinpoint chemical modification study of Na^+^-NQR, we previously proposed that the N-terminal region of NqrB is oriented toward the membrane phase rather than protruding into the cytoplasmic medium along the surface of NqrA (*21*), as implied by the crystallographic structure (*16*). The present cryo-EM structure unambiguously corroborates this view. We note that the three cryo-EM structures of Na^+^-NQR (i.e. the enzyme without bound inhibitor and the enzymes with bound korormicin A or aurachin D-42) are almost identical except for the N-terminal region of NqrB, which is disordered in the absence of inhibitor (Extended Data Figure 6A and 6C).

**Figure 6:**
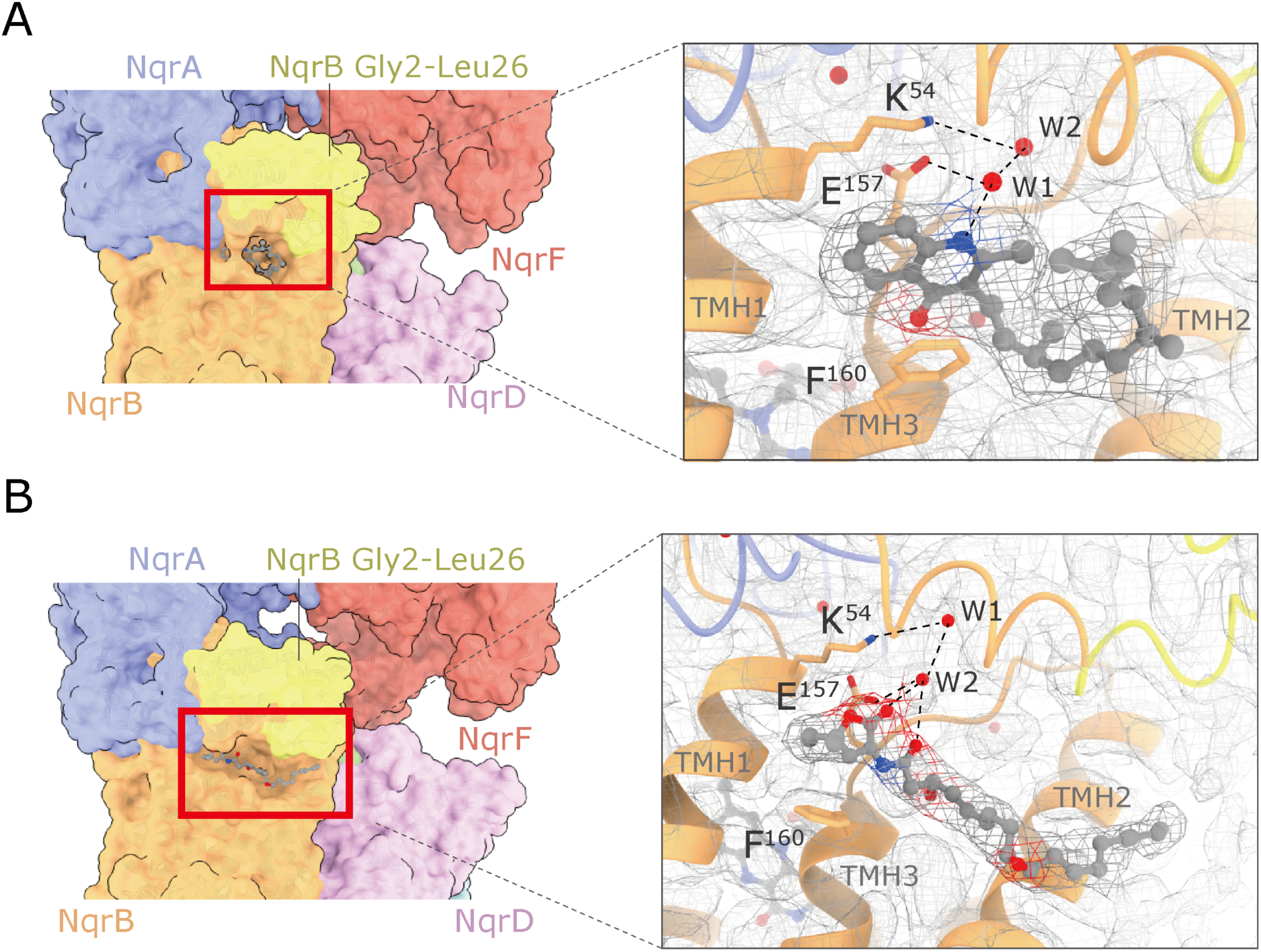
Structures of Na^+^-NQR with bound inhibitors. Aurachin D-42 (**A**) and korormicin A (**B**) bind to the N-terminal region starting with TMH 1 of NqrB. *Left* and *right* panels show the overall and enlarged views of the binding pockets, respectively. The residues that are involved in the interaction with the inhibitors are indicated as *sticks* model. The density maps are shown as *mesh* representations. The disordered region (NqrB-Gly2–Leu26) in the absence of inhibitor is indicated in *yellow*.

The binding model of aurachin D-42 (a naphthoquinone-like inhibitor) is shown in Figure 6A and Extended Data Figure 7A. The NH group on the quinolone ring forms a hydrogen bond with NqrB-Glu157 via an intervening water molecule. The configuration of NqrB-Glu157 may be fixed by a hydrogen bond with the adjacent NqrB-Lys54 via a water molecule. NqrB-Phe160 is involved in a π-stacking interaction with the quinolone ring, which may lock aurachin D-42 inside the binding cavity (Extended Data Figure 7A). The farnesyl chain bends at the second isoprene unit and turns toward the membrane phase to avoid steric obstruction arising from the cavity wall formed by TMH 2.

The binding model of korormicin A is presented in Figure 6B and Extended Data Figure 7B. The two carbonyl groups, one is on the lactone ring and the other is in the amide moiety, form hydrogen bonds with NqrB-Glu157 via an intervening water molecule. The configuration of the lactone ring relative to the adjacent amide plane appears to be restricted by steric obstruction arising from Met57 and Ile58 on TMH 1 and Phe160 on TMH 3 of NqrB (Extended Data Figure 7B). The 3’-OH points into the cavity interior. Although the amino acid residue(s) that can make a hydrogen bond to the OH group cannot be definitely identified, this functional group appears to orient toward the amide bond connecting NqrB-Glu157 and -Gly158 on TMH 3. The long alkyl side chain extends toward a hydrophobic cleft composed of Trp23, Leu26, Phe142, and Val145 of NqrB and is buried in it. No hydrogen-bond donating residue was identified near the oxygen atom of the epoxy group at the 9’/10’ positions, which orients toward the outside of the cavity. In contrast to the aurachin D-42-bound structure, NqrB-F160 does not face toward korormicin A.

The above results strongly suggest that NqrB-Glu157 plays a key role in the binding of the two inhibitors by making a hydrogen bond to their toxophoric moieties. To assess the significant contribution of NqrB-Glu157, we prepared a NqrB-Glu157Ala mutant and compared the inhibitory activities of korormicin A and aurachin D-42 to those measured with the wild-type enzyme. The NADH-UQ_1_ oxidoreductase activity of the isolated NqrB-Glu157Ala enzyme without inhibitor is 50–60% lower than that of the wild-type enzyme. This is probably because the binding site of UQ in NqrA, which may be located near the inhibitor binding site (*18*), is influenced by the mutation. While IC_50_ values of korormicin A and aurachin D-42 determined in the NADH-UQ_1_ oxidoreductase assay with the wild-type enzyme were one-digit nM levels (*18, 19*), 50% inhibition was not observed for either inhibitor with the mutant, even at 2.0 µM. These results indicate the critical contribution of NqrB-Glu157 to the inhibitor binding.

We previously sought a new method to enable pinpoint chemical modification of amino acid residue located in the N-terminal region of NqrB. We found that *N*-acyl-*N*-alkyl sulfonamide chemistry (Supplementary Figure 2), one of the protein-ligand affinity-driven substitution techniques (*27*), to be suitable for this purpose. Using this method, an electrophilic *N*-acyl-*N*-alkyl sulfonamide group attached to the end of the side chain of korormicin derivatives (NAS-K1 and NAS-K2, Supplementary Figure 2) specifically reacts with the nucleophilic NqrB-Lys22 (*21*). This result strongly suggests that NqrB-Lys22 is located near the side chain of the korormicin derivatives. In corroboration of this, in the cryo-EM structure, NqrB-Lys22 is located in the vicinity of the hydrophobic cleft that accommodates the side chain of korormicin (Extended Data Figure 7B), although the position of the side chains of NAS-K1 and -K2 would not necessarily be identical to that of natural korormicin A because of their structural differences.

## DISCUSSION

A complete picture of the electron transfer reactions and the mechanism of Na^+^ translocation coupled to electron transfer in Na^+^-NQR remains elusive. High-resolution structures of Na^+^-NQR are essential for understanding these central questions. Here, we describe the cryo-EM structures of *V. cholerae* Na^+^-NQR with and without a bound inhibitor at 2.5- to 3.1-Å resolution. The present study provides new information about the enzyme structure, which sheds light on unsettled issues in the earlier X-ray crystallographic study (*16*). First, we revealed how the cytoplasmic water-soluble domains of the NqrA and NqrF subunits attach to the membrane embedded subunits. The C-terminal region of NqrA (Arg330−Asn379) is intertwined with the protruding part of the N-terminus of NqrB (Phe34−Leu53) (Extended Data Figure 5). The soluble domain of NqrF is anchored to the soluble domain of NqrA by tight electrostatic interactions (Figure 2), which may allow the rest of NqrF to flexibly move. Second, riboflavin is located inside NqrB at the cytoplasmic side, where it is surrounded by central TMHs 1, 3, 5, and 8 (Figure 4). Therefore, it is not conceivable that the Na^+^ channel traverses the center of NqrB (*16*). Third, the unique density between NqrD and NqrE can be assigned to a 2Fe-2S cofactor (not a single iron atom), which is coordinated by the two pairs of the conserved cysteine residues (NqrD-Cys29/NqrE-Cys120 and NqrE-Cys28/NqrD-Cys112) (Figure 3). Finally, we found that both korormicin A and aurachin D-42 bind to the N-terminal region of NqrB (Figure 6), corroborating the results of our previous photoaffinity labeling experiments (*18, 19*). It is noteworthy that while the N-terminal region of NqrB (Gly2−Leu26) is disordered in the absence of inhibitors, the binding of inhibitors gives rise to a distinct conformation that we could identify by cryo-EM. Altogether, the present cryo-EM study provides a definite foundation for understanding the function of Na^+^-NQR and the molecular mechanism of its specific inhibitors. To elucidate the predicted large structural changes during the catalytic turnover, individual cryo-EM structures of different partially-reduced states of the enzyme are necessary.

The UQ reduction is the final step in the overall electron transfer pathway in Na^+^-NQR. We recently found that the role of UQ is not simply to accept electrons from the reduced riboflavin^NqrB^ to reset the catalytic cycle but critically regulates Na^+^-translocation via its side chain moiety (*28*). Therefore, identification of the UQ reaction site is essential for fully understanding of the overall electron transfer and the mechanism responsible for Na^+^-pumping driven by the electron transfer. It has been suggested, in some literature (*10, 26, 29*), that the isolated Na^+^-NQR contains sub-stoichiometric amounts of a high affinity bound UQ_8_; however, the present cryo-EM structure presents no evidence for the existence of the bound UQ_8_. Photoaffinity labeling experiments using a photoreactive UQ showed that the UQ head-ring binds to the contact area of NqrA-Leu32−Met39 and Phe131−Lys138 (*18*). In the current cryo-EM structure of an oxidized form of Na^+^-NQR, this area forms a part of a cavity that is large enough to accommodate the UQ head-ring (Extended Data Figure 8), but the cavity is ∼20 Å above the predicted membrane surface. Therefore, the distance between the cavity and riboflavin located in NqrB is too long (∼30 Å) to allow efficient electron transfer. Unfortunately, we have yet to obtain the cryo-EM structure of Na^+^-NQR with bound UQ_8_ and/or short-chain UQs such as UQ_2_. This is presumably because the binding affinities of substrate UQs to the enzyme are substantially lower compared to the potent inhibitors used in the present study. This challenging task is under way in our laboratory.

Steuber et al. (*16*) proposed that a membrane spanning narrow cavity formed by the central TMHs 1, 3, 6, and 8 in NqrB is the Na^+^-channel, in which Na^+^ travels from the cytoplasmic to periplasmic sides. Based on the current cryo-EM structure, this possibility may be low because riboflavin lies at the center of NqrB on cytoplasmic side, where it is surrounded by TMHs 1, 3, 5 and 8 (Figure 4) and because the membrane interior loop (Gly266–Ser276), whose position is substantially different from that in the crystallographic structure (*16*), covers the cavity interior facing the periplasmic side (Figure 1C). Thus, there is no membrane spanning cavity in the center of NqrB. Considering that Na^+^-NQR transports Na^+^ against the electrochemical gradient, it is not conceivable that a channel that completely traverses the membrane could exist in any active conformation of the enzyme.

Korormicin A is a highly specific and potent inhibitor of Na^+^-NQR, which exhibits no inhibition against the respiratory enzymes in mammalian mitochondria (*3, 20*). To identify important structural factors of korormicin A required for exhibiting this inhibition, we previously investigated the relationship between structure and inhibitory potency using a series of synthetic korormicin A analogs (*19*), as summarized in Supplementary Figure 3. The cryo-EM structure of the korormicin A-bound Na^+^-NQR allows us to understand how such structural factors are involved in the interaction with the enzyme. Here, we focus the following three important structural factors: i) the alkyl branches (CH_3_/C_2_H_5_) at the 5-position, ii) *R*-configuration of the 3’-OH group, and iii) the epoxy group at the 9’/10’ positions.

1. The presence of the alkyl branches at the 5-position of the lactone ring is essential for the potent inhibition, as seen from the drastic decrease in the inhibitory potency of compound D3 (Supplementary Figure 3). In unbound korormicin A, the lactone ring can freely rotate around the bond connecting it to the amide (-NH-CO-) moiety and, hence, does not have a fixed conformation relative to the amide plane. However, in the bound state, steric obstruction between the alkyl branches and the cavity wall formed by NqrB-Met57 and - Ile58 on TMH 1, NqrB-Phe160 on the TMH 3, and NqrA-Try337 (Extended Data Figure 7B) may fix the conformation of the lactone ring and make its carbonyl group orient toward the inside of the binding cavity, enabling a hydrogen bond to NqrB-E157 via intervening water molecules (Figure 6B).
2. The inhibitory potency of the 3’-epimer of korormicin A markedly decreased (korormicin A vs. compound D4), indicating the importance of the *R*-configuration of the OH group at the 3’-position. This functional group appears to orient toward the amide bond connecting NqrB-Glu157 and -Gly158 on TMH 3 to form a hydrogen bond (Figure 6B and Extended Data Figure 7B). In contrast, the OH group in *S*-configuration of compound D4 may not orient toward the inside of the binding cavity.
3. The epoxy group at the 9’/10’ positions is favorable but not essential for inhibition (korormicin A vs. compound D5). Since no hydrogen-bond-donating residue is located near the epoxy oxygen atom, which orients toward the outside of the binding cavity (Figure 6B), this favorable effect may not be due to a hydrogen bond. It is rather likely that the presence of the epoxy group adjacent to the 4’,6’-diene plane bends the alkyl side chain moiety, which enables the side chain to extend toward the hydrophobic cleft composed of NqrB-Trp23, -Leu26, -Phe142, and -Val145 by making a detour around the cavity wall formed by TMH 2 (Figures 6B and Extended Data Figure 7B).

The molecular interactions between aurachin D-42 and the binding cavity are slightly different from those for korormicin A described here (Figure 6A vs 6B), suggesting that the N-terminal region of NqrB flexibly enfolds the inhibitors according to their unique chemical frameworks. Our cryo-EM structures of the inhibitor-bound Na^+^-NQR can help molecular design strategies for new inhibitors targeting the enzyme.

Finally, we discuss a unique phenomenon observed in the previous photoaffinity labeling study using photoreactive aurachin D-type inhibitors ([^125^I]PAD-1 and [^125^I]PAD-2, Supplementary Figure 1) with the isolated *V*. *cholerae* Na^+^-NQR (*18*). The labeling by these [^125^I]-incorporated inhibitors rather than being competitively suppressed in the presence of excess other inhibitors (including their non-radioactive analogs PAD-1 and PAD-2), was *enhanced* when a molar ratio of ^125^I-incorporated inhibitor to the enzyme is relatively low (<10). This unusual competitive behavior is difficult to reconcile with a simple scenario in which different inhibitors share a common binding site. To explain this, we proposed an equilibrium model for the binding of ^125^I-incorporated inhibitor and competitor based on the assumption that there are two distinct inhibitor-bound states (i.e. one-inhibitor- and two-inhibitor-bound states), in which the yields of the labeling reaction are considerably different (ref. *18* for details). While we cannot exclude other scenarios that would explain the unusual competitive behavior, this model accounted for the consecutive changes in the nature of the competition (from enhancement to suppression) as the concentration of competitor increases. However, the present cryo-EM structures provided no evidence for the existence of two distinct binding sites for inhibitor; therefore, the equilibrium model must be corrected.

The current study revealed that, while the N-terminal region (Gly2−Leu26) of NqrB is disordered in the absence of inhibitor, the binding of inhibitor gives rise to a distinct conformation of this region. It is, therefore, likely that the bound inhibitor shapes the region into its own binding cavity reflecting its chemical framework. This mode of binding may be thought of as the so-called “induced-fit binding”. Given that the inhibitor binding is a reversible equilibrium process, the binding cavity of the inhibitor may exist in, at least, two different conformations in equilibrium: one, disordered and indefinite conformation with a low affinity to inhibitor and the other, distinct fixed conformation with a high affinity to inhibitor. The latter would correspond to the binding cavity identified by cryo-EM (Figure 6A and 6B). Based on this idea (namely, two different conformations of a single binding cavity), we undertook to produce a new equilibrium model to explain the unusual competitive behavior between ^125^I-incorporated inhibitor and competitor (Supplementary Figure 4A in Appendix Discussion). Note that although there could be many consecutive intermediates between the two extreme conformations, to simplify the analysis we have assumed only two conformations. We estimated the concentration of the enzyme-^125^I-incorporated inhibitor complex (“F-[^125^I]I” complex in Supplementary Figure 4A), to which the radioactivity incorporated into the enzyme by photolysis is proportional, under varying modeling conditions. The details of this analysis are described in Supplemental Discussion. In conclusion, although the analytical results change depending on the parameters (e.g. relative magnitudes of the equilibrium constants *K*_1_–*K*_4_), the concentrations of the F-[^125^I]I complex change, from enhancement to suppression, as the concentration of competitor increases in most modeling sets. This is because the added competitor, in a certain concentration range, increases the proportion of the binding cavity in the fixed conformation by reversibly binding to the enzyme. Thus, the unusual competitive behavior observed in the previous photoaffinity labeling studies (*18*) can be accounted for by an equilibrium model based on the idea of two different conformations of a single binding cavity. The current structural study led to this new equilibrium model, both by ruling out the presence of two distinct binding sites, and suggesting the possibility of multiple conformations of a single binding site with different affinities.

## METHODS

### Expression of wild-type and mutant Na^+^-NQRs

The recombinant Na^+^-NQR of *V. cholerae*, which contains a six-histidine affinity tag at the C-terminus of NqrF subunit, was produced in a *V. cholera* strain lacking the genomic *nqr* operon (*4*). Δ*nqr* cells containing the wild-type or mutant *nqr* operon on a pBAD plasmid were grown in Luria-Bertani (Miller) medium in 30-liter fermenters (New Brunswick BF-5000, Microbiology Core Facilty, CBIS, RPI) at 37 °C with constant agitation (300 rpm) and aeration of 20 L/min. Expression of the *nqr* operon was induced by arabinose. Cells were harvested, washed by centrifugation, and disrupted by passing through a Microfluidizer at a pressure of 20,000 p.s.i. (*30*). The membrane preparations were obtained by ultracentrifugation (100,000 g) and washed with a buffer containing 50 mM NaPi, 5.0 mM imidazole, 300 mM NaCl, and 0.05% glycerol. The site-directed mutant NqrB-Glu157Ala was obtained using the QuickChange II XL mutagenesis kit (Agilent), as reported before (*30*). The forward primer was 5’-CGCAAGCATGAAGTCAACGCTGGTTTCTTCGTTACCTCTAT-3’ and the reverse primer was complementary to the forward primer.

### Purification of Na^+^-NQR

The membrane pellets (40 g) were solubilized in 600 mL of solubilization buffer [50 mM NaPi, 300 mM NaCl, 5.0 mM imidazole, and protease inhibitor cocktail (Sigma-Aldrich), pH 8.0] containing 0.4% (w/v) *n*-dodecyl-β-D-maltoside (DDM) with stirring for 1 h at 4 °C (*4*). The mixture was clarified by ultracentrifugation (100,000 g), and supernatant was mixed with 20 mL of Ni-NTA resin (Qiagen) that had been equilibrated with binding buffer (50 mM NaPi, 300 mM NaCl, 5.0 mM imidazole, and 0.05% DDM). After gentle agitation for 1 h at 4 °C, the suspension was poured into an empty plastic column. The column was washed with 4 volumes of binding buffer, followed by 5 volumes of wash buffer (same as binding buffer but containing 10 mM imidazole). The protein was eluted from the column using elution buffer (same as binding buffer but containing 100 mM imidazole).

The resulting Na^+^-NQR preparation was applied to a DEAE anion exchange column (HiPrep DEAE FF 16/10, connected to an ÄKTA system, Cytiva) equilibrated with buffer A [50mM Tris/HCl, 1.0 mM EDTA, 5% (v/v) glycerol, and 0.05% (w/v) DDM, pH 8.0]. The protein was eluted with a linear gradient of buffer B (buffer A containing 2.0 M NaCl), and the fractions containing pure Na^+^-NQR were pooled and concentrated 15∼20 mg of protein/mL. Subunit assembly was verified by SDS-PAGE (*31*) and enzyme activity was checked by the inhibitor-sensitive NADH-UQ_1_ oxidoreductase activity measurements at 282 nm (*ε* = 14.5 mM^- 1^cm^-1^, *18*) (Extended Data Figure 3).

For the preparation of EM grids, an aliquot of this enzyme preparation (50 µL, 780 µg) was further purified by chromatography on a Superose^TM^ 6 Increase 10/300 gel filtration column (Cytiva), equilibrated with buffer (50 mM Tris-HCl, 100 mM NaCl, 1.0 mM EDTA, 0.05% DDM, pH 8.0) to remove glycerol from the sample. The peak fraction containing Na^+^-NQR (3.0 mL) was concentrated to 20 µL using an Amicon Ultra 100 K centrifugal filter (Merck-Millipore), and used immediately for the cryo-EM grid preparation.

### Cryo-EM acquisition

To prepare the cryo-EM grid for purified Na^+^-NQR, 2.7 μL of the enzyme solution (50 µM) was loaded onto a glow-discharged Quantifoil Cu R1.2/1.3 and blotted by Vitrobot IV followed by vitrification with liquid ethane. To obtain samples with bound aurachin D-42 (abbreviate as “Na^+^-NQR^AD42^”) or korormicin A (abbreviate as “Na^+^-NQR^KA^”), the enzyme and inhibitor were mixed at a 1:10 molar ratio and incubated for 10 seconds before loading on a grid. Cryo-EM movies were automatically acquired using a Titan Krios electron microscopy (Thermo Fisher, USA) equipped with K3 BioQuantum camera (Gatan, United States) using SerialEM software (*32*). The cryo-EM movies were collected at a nominal magnification of 81,000 and the pixel size was 0.88 Å/pix. The total electron dose and frame rate were 65 electrons/Å^2^ and 0.1 s, respectively.

### Image processing

Image processing steps for each condition are summarized in Extended Data Figure 1. All steps were performed by cryoSPARC v3.2.0 or v3.3.1 software (*33*). Topaz software was used for machine-learning based particle picking (*34*). For native Na^+^-NQR, Na^+^-NQR^AD42^, and Na^+^- NQR^KA^, 6,336, 11,430, and 10,406 movies were used, respectively. After motion correction and constant transfer function (CTF estimation), auto-picking based on Topaz machine-learning resulted in 634,154, 3,350.484, and 721,916 particles for Na^+^-NQR, Na^+^-NQR^AD42^, and Na^+^-NQR^KA^, respectively. Particles were extracted from the motion-corrected micrographs and selected using Heterogenous Refinement. The selected particles were subjected to Non-uniform Refinement with the optimizing per-particle defocus option. The refinement provided density maps for Na^+^-NQR, Na^+^-NQR^AD42^, and Na^+^-NQR^KA^ at 2.7-, 2.5-, and 2.6-Å resolution, respectively.

The density corresponding to the cytoplasmic domain of NqrF was weak and blurry due to its flexibility. To obtain the structure of this domain, local refinement on NqrA and NqrB was carried out after Non-uniform Refinement. Then, focused 3D classification on NqrF and NqrA without alignment was applied to the results of local refinement. Of the results of focused 3D classification, the particles belonging to the density maps, in which NqrF was clearly observed, were selected: three classes for Na^+^-NQR (Class 1; 82,790 particles, Class 2; 80,882 particles, Class 3; 72,234 particles), one class for Na^+^-NQR^AD42^ (55,375 particles), and one class for Na^+^-NQR^KA^ (50,444 particles). The selected particles were subjected to Non-uniform Refinement. As a result of these procedures, three density maps of Na^+^-NQR without bound inhibitor, in which the soluble domain of NqrF was clearly defined, were obtained, all at 3.1-Å resolution. For Na^+^-NQR^AD42^ and Na^+^-NQR^KA^, density maps at 3.0-Å and 2.9-Å resolution, respectively, were obtained. Resolution was based on the gold standard criterium: Fourier shell correlation = 0.142 criterion.

### Model building and refinement

To build the atomic model of Na^+^-NQR, each subunit modeled in the crystallographic structure (PDBID: 4P6V) was fitted into the density map as a rigid body. The initial model was flexibly fitted against the map using the Chain Refine function of COOT software (*35*). Then, the roughly fixed model was extensively manually corrected, residue by residue, using the Real Space Refine Zone function in COOT. The manually corrected model was refined using the phenix_real_space_refine program (*36*) with secondary structure and Ramachandran restraints. The resulting model was manually checked by COOT. The water molecules were placed in the models manually. This iterative process was performed for several rounds to correct remaining errors until the model satisfied the geometry. The model quality was assessed by Molplobity and EMRinger scores (*37, 38*). For model validation against over-fitting, the final models were used to calculate Fourier shell correlation against the final density maps employed in model building using the phenix refine program. The statistics of maps and models presented here are summarized in Supplementary Tables 1 and 2. All figures were prepared using UCSF chimeraX software (*39*).

### Data availability

The cryo-EM maps have been deposited in the EMDB under accession codes, 33242, 33243, 33244, 33245, and 33246. The consensus maps for Na^+^-NQR, Na+-NQR^AD42^, and Na^+^-NQR^KA^ were also deposited as additional maps in the depositions, 33242, 33245, and 33246. The atomic models have been deposited in the Protein Data Bank under accession codes, 7XK3, 7XK4, 7XK5, 7XK6, and 7XK7. The initial model for model building is accessible in PDB under accession number 4P6V. The data that support the findings of this study are available from the corresponding author upon reasonable request.

## Acknowledgement

We thank Dr. Joel E. Morgan (Rensselaer Polytechnic Institute) for helpful discussion. This work was supported by JSPS KAKENHI (Grant Numbers JP21H02130 to H.M., 22K14837 to T.M., 20K06514 to J.K., and 22H02273 to M.M.), and the National Institutes of Health (RO1-AI132580 to B.B.).

## Author contributions

J.K., M.I., T.M., M.M., B.B., and H.M. designed the research; M.I., T.M., N.L.B., and B.B. expressed and purified Na^+^-NQR; J.K., M.I., and T.M. performed EM sample preparation and data collection. J.K., M.I., and T.K. performed structural determination and built the atomic models. Y.K., and M.I. analyzed the equilibrium model. J.K., M.I., T.M., M.M., B.B., and H.M. analyzed data; H.M. directed the project and wrote the paper with J.K., I.M., T.M., M.M., and B.B.

## Extended Data Figures

**Extended Data Figure 1:**
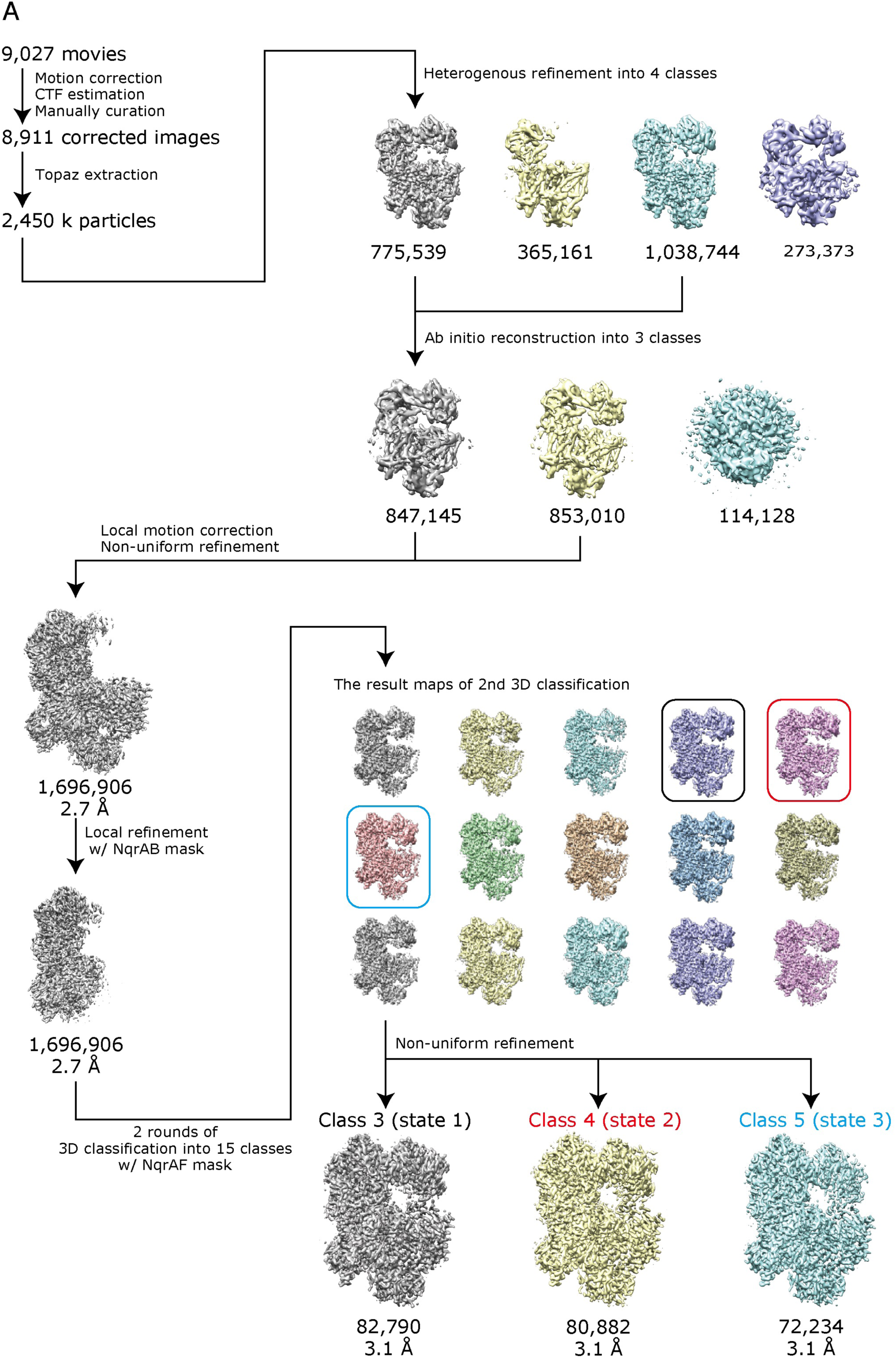
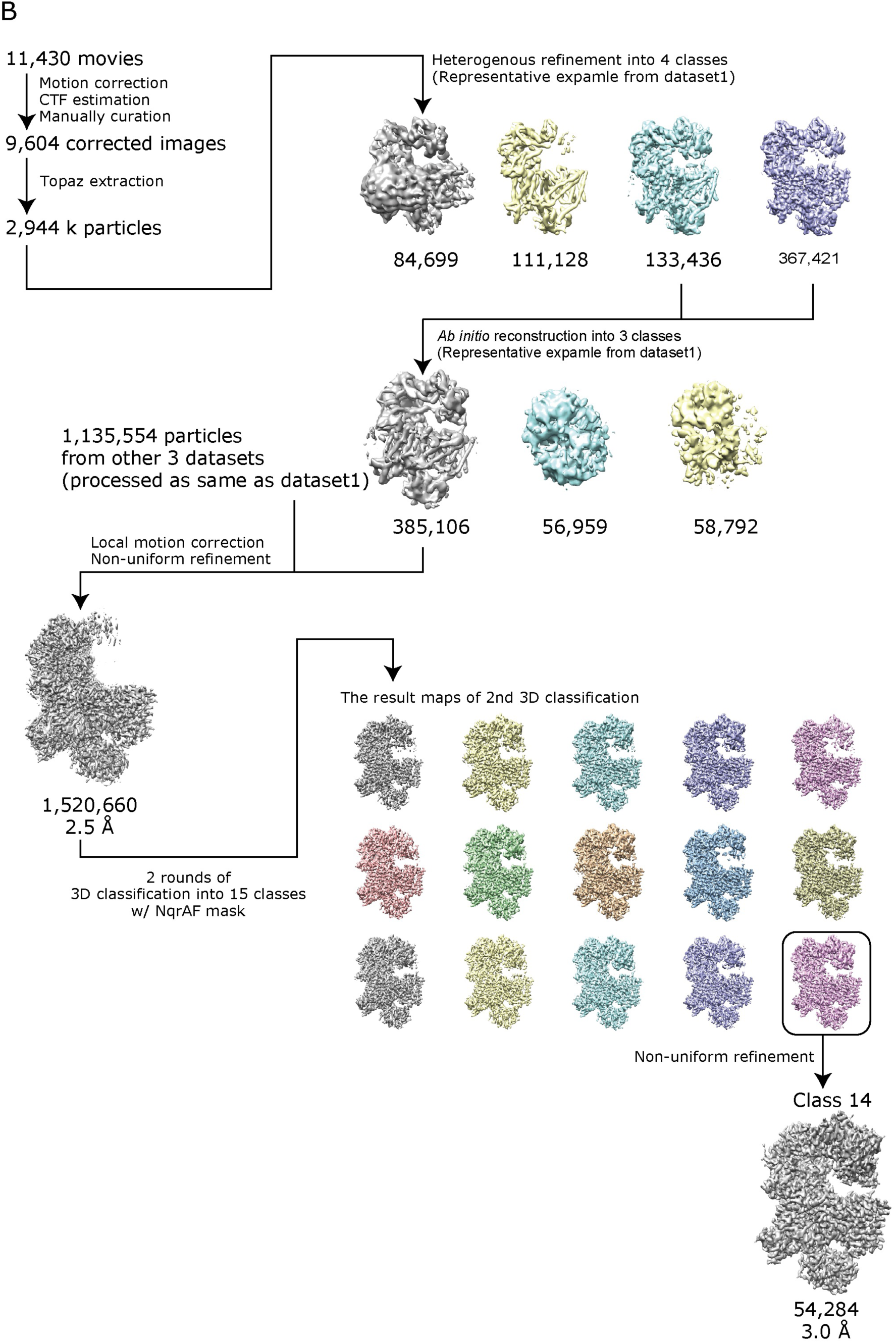
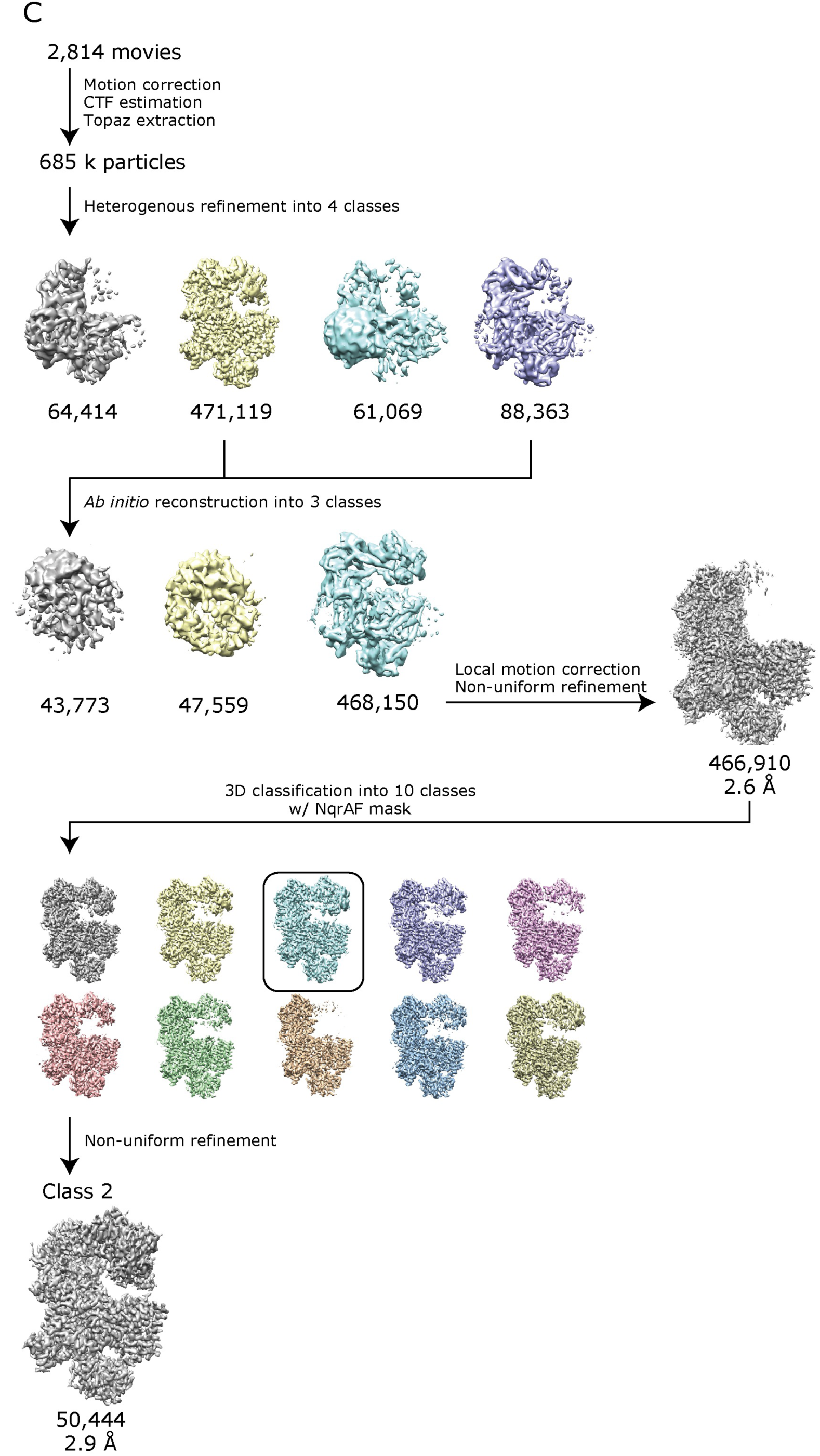
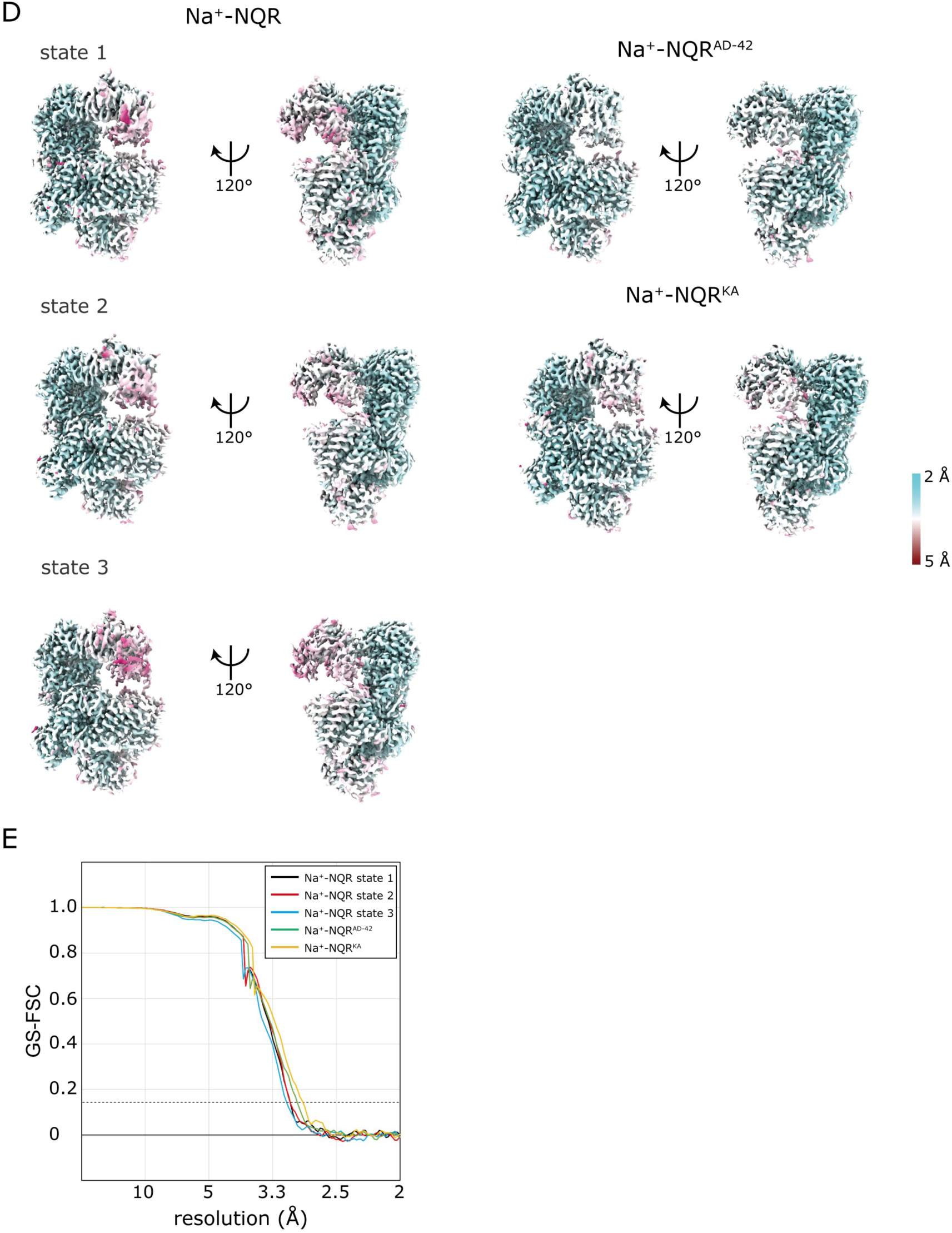
Flow charts for image processing of three Na^+^-NQR preparations. (**A**) Na^+^-NQR without inhibitor, (**B**) Na^+^-NQR with bound aurachin D-42 (Na^+^-NQR^AD42^), and (**C**) Na^+^-NQR with bound korormicin A (Na^+^-NQR^KA^). For the dataset of Na^+^-NQR^AD42^ (total 11,430 movies), the dataset was separated into four set. The image processing of each set was conducted according to the flow chart. Representative results of the four dataset is shown in a panel **C**. The selected particles from each dataset were merged before Homogenous refinement. (**D**) Cryo-EM density maps of Na^+^-NQR (states 1–3), Na^+^-NQR^AD42^, and Na^+^-NQR^KA^. The maps are colored according to local resolution as indicated in the color bar. (**E**) Gold-standard Fourier shell correlation (GS-FSC) curves for each Na^+^-NQR, using GS-FSC = 0.142 for resolution criterion (dotted line).

**Extended Data Figure 2:**
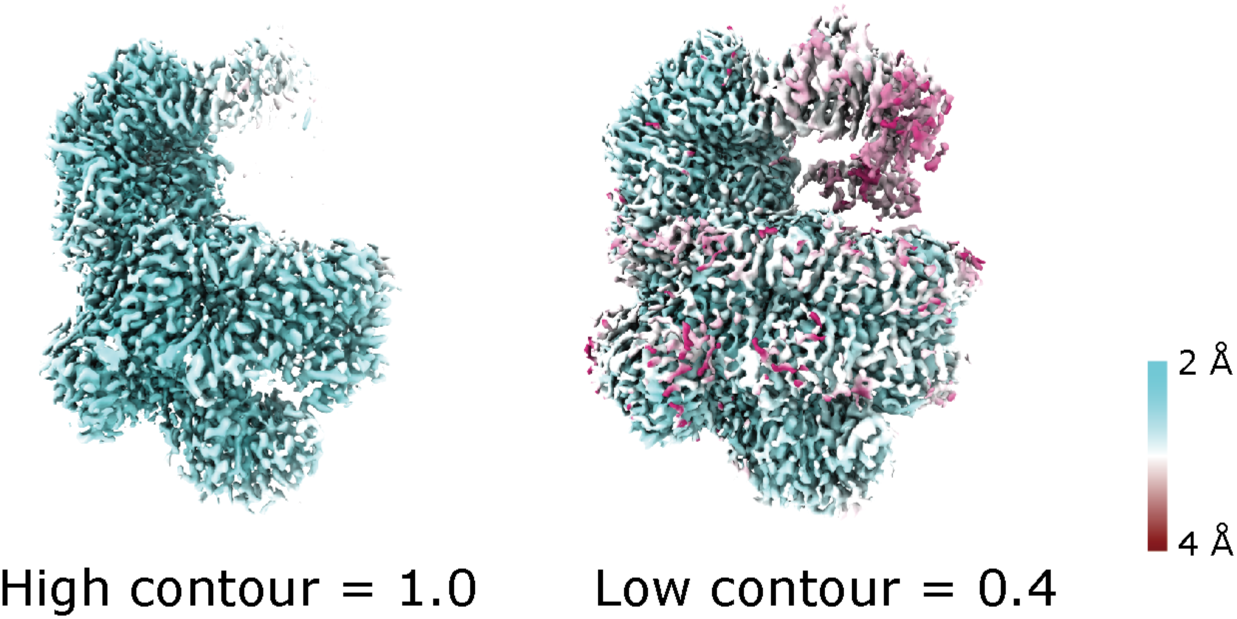
The consensus map for Na^+^-NQR. The consensus map for Na^+^-NQR is represented as solid surfaces. The map was obtained Non-Uniform refinement using all selected particles (**Extended Data Figure 1**). The maps are colored by local resolution indicated as color key. The contour levels are set to high (*left*) and low (*right*). The density corresponding to the hydrophilic domain of NqrF are weak and has low resolution compared to other subunits.

**Extended Data Figure 3:**
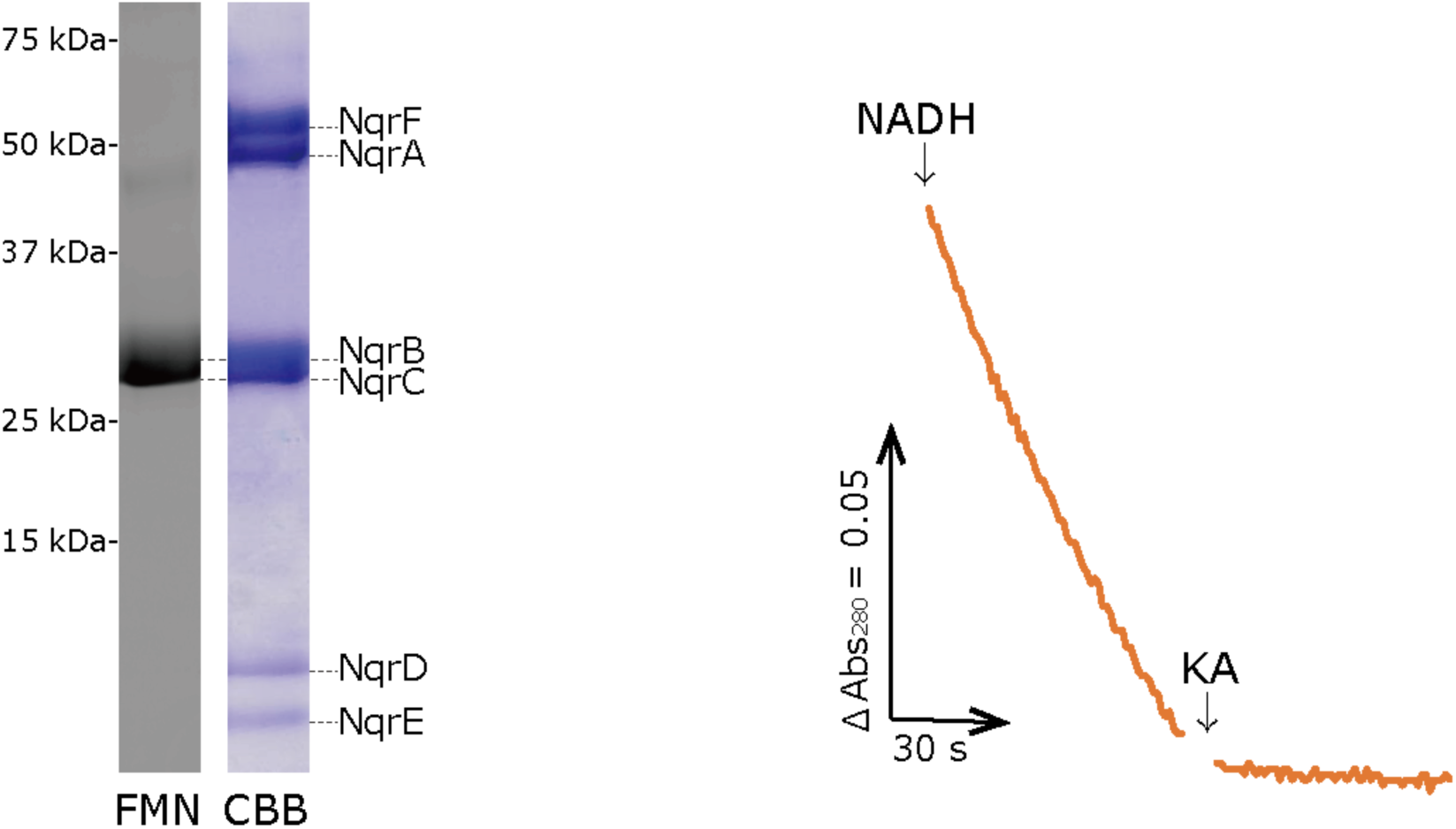
Biochemical characterization of the purified Na^+^-NQR. SDS-PAGE of the purified *V. cholerae* Na^+^-NQR. The subunits were stained with CBB R-250. FMNs in the NqrB and NqrC subunits were visualized using bio-imaging analyze Typhoon FLA-9500 (Cytiva) using a 473 nm light source and an LPB emission filter (emission wavelengths shorter than 510 nm are cut off). The NADH-UQ1 oxidoreductase activity of Na^+^-NQR was almost completely inhibited by korormicin A (1.0 µM).

**Extended Data Figure 4:**
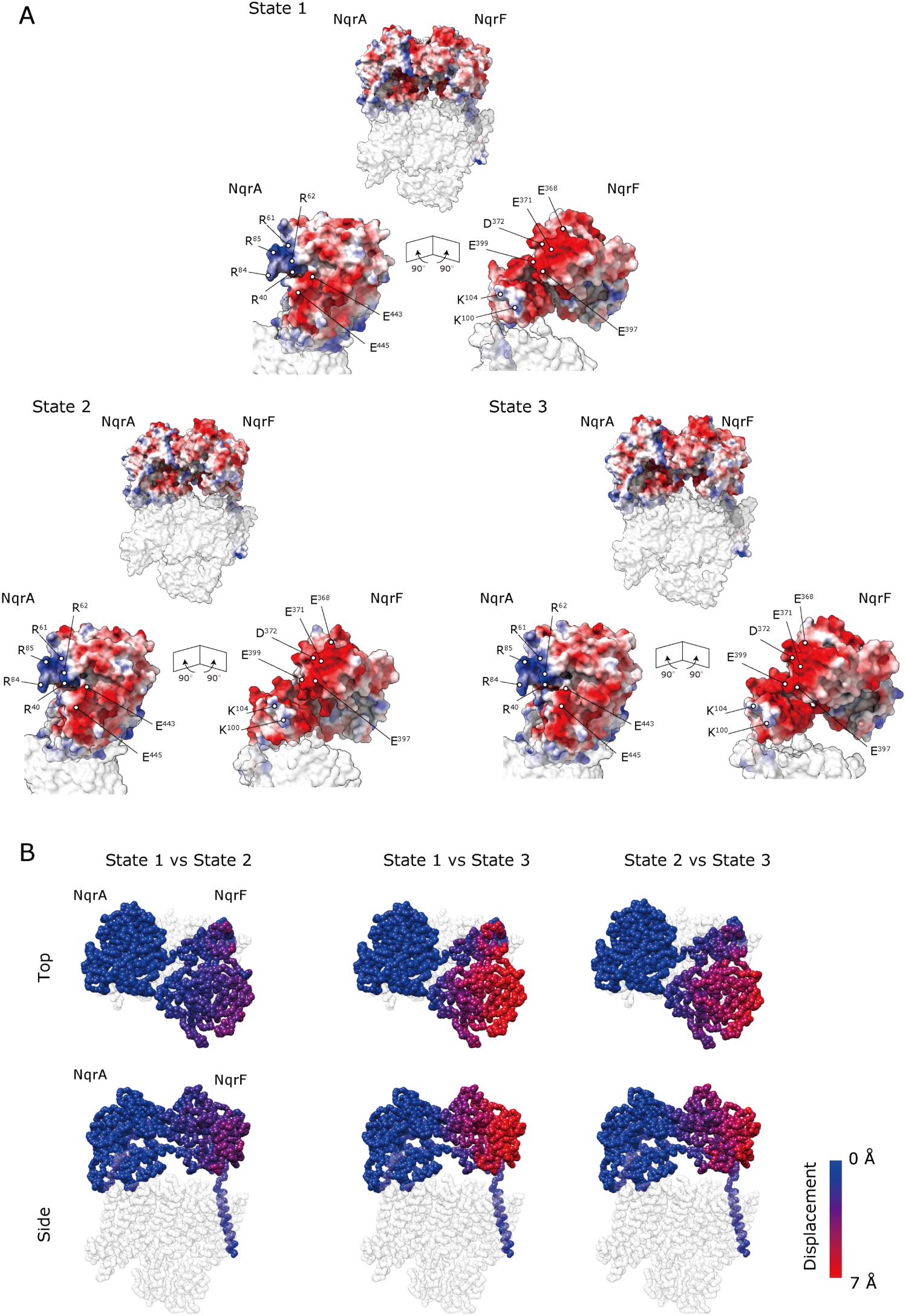
Electrostatic interactions between NqrF and NqrA in the 3 states. (**A**) The structures of the interacting area between NqrF and NqrA in states 1–3. The structures of the interacting area are almost identical one another irrespective of significant structural differences of the rest of the hydrophilic parts of NqrF. The positive patch formed by R40, K61, K62, R84, and R85 of NqrA interacts with the negative patch formed by E368, E371, E372, E397, and E399 of NqrF in the three states. Negative tips of NqrA (E443 and E445) are in close proximity to positive tips of NqrF (K100 and R104). (**B**) Each state of Na^+^-NQR was superimposed on the NqrB subunit. NqrA and NqrF are represented as a sphere model and colored by displacement values between the three states calculated for the Cα atoms; *blue* (small changes) to *red* (large changes). Other subunits are shown in semi-transparent.

**Extended Data Figure 5:**
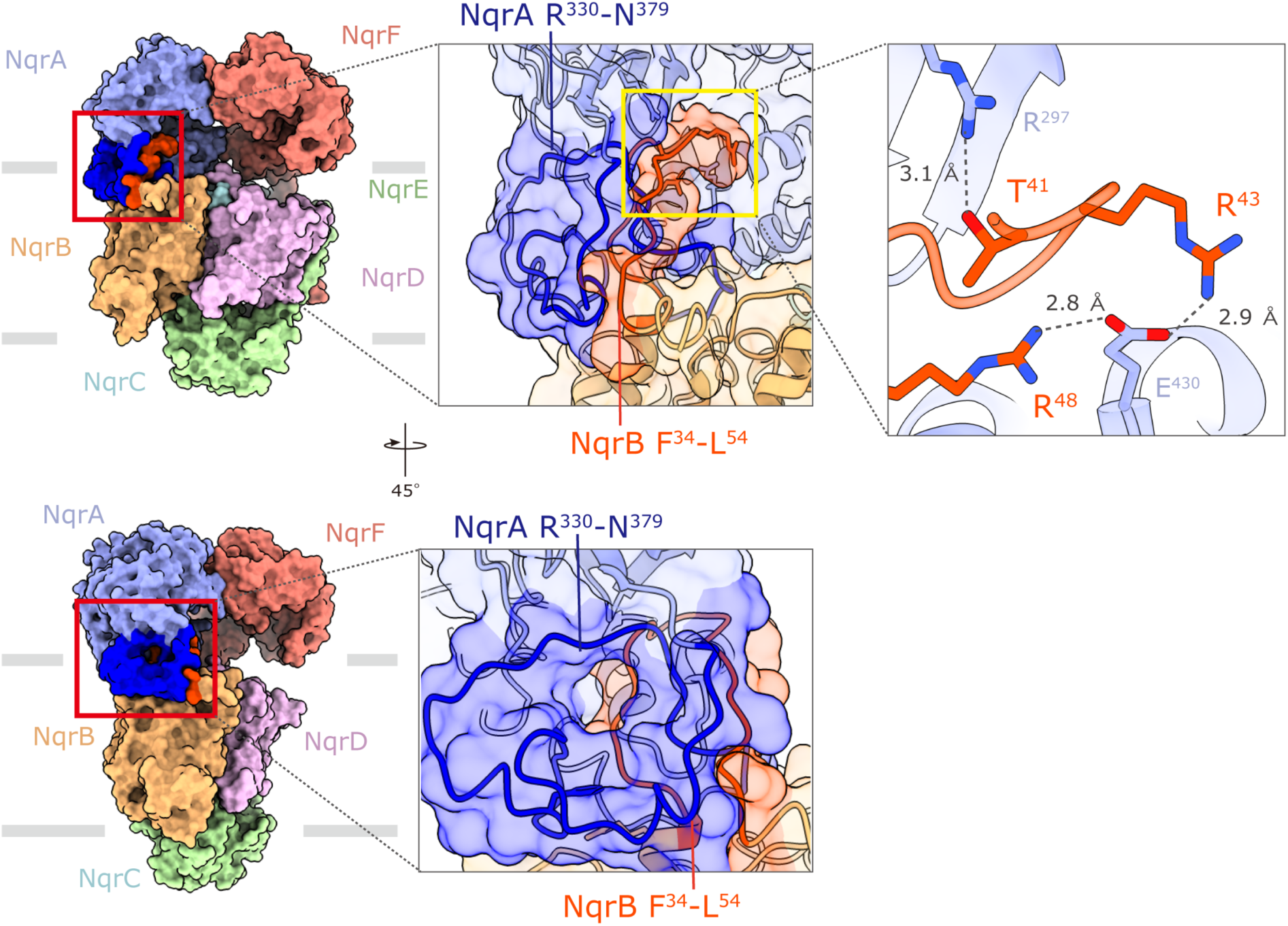
Structure of the contact area between NqrA and NqrB. The contact area between the C-terminal region of NqrA (Arg330−Asn379, in *dark blue*) and the protruding part of the N-terminal region of NqrB (Phe34−Leu54, in *red*).

**Extended Data Figure 6:**
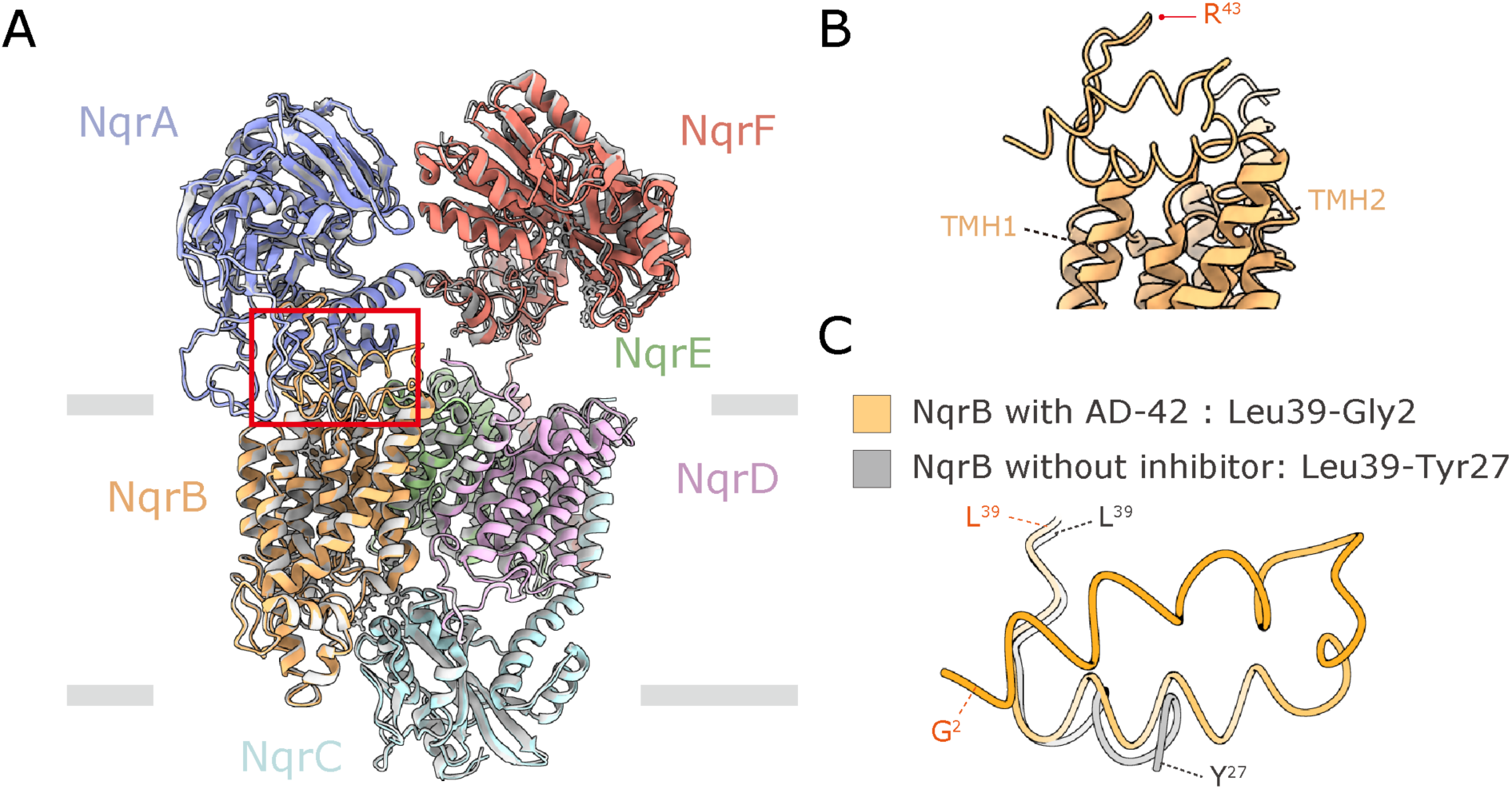
Comparison of the Na^+^-NQR structures between aurachin D-42-bound and non-bound states. (**A**) Overlay of Na^+^-NQR structure with bound aurachin D-42 (color) with the structure in the absence of inhibitor (gray). The two structures are almost identical except for the N-terminal region of NqrB indicated by a red square. (**B**) The structure of the protruding N-terminal stretch starting with TMH 1 of NqrB in Na^+^-NQR with bound aurachin D-42. The N-terminal stretch first protrudes from the membrane phase and then turns back toward the membrane by bending at NqrB-Arg43. (**C**) Close-up view of the N-terminal region of NqrB. The region Gly2–Leu26 is disordered in the absence of the inhibitor.

**Extended Data Figure 7:**
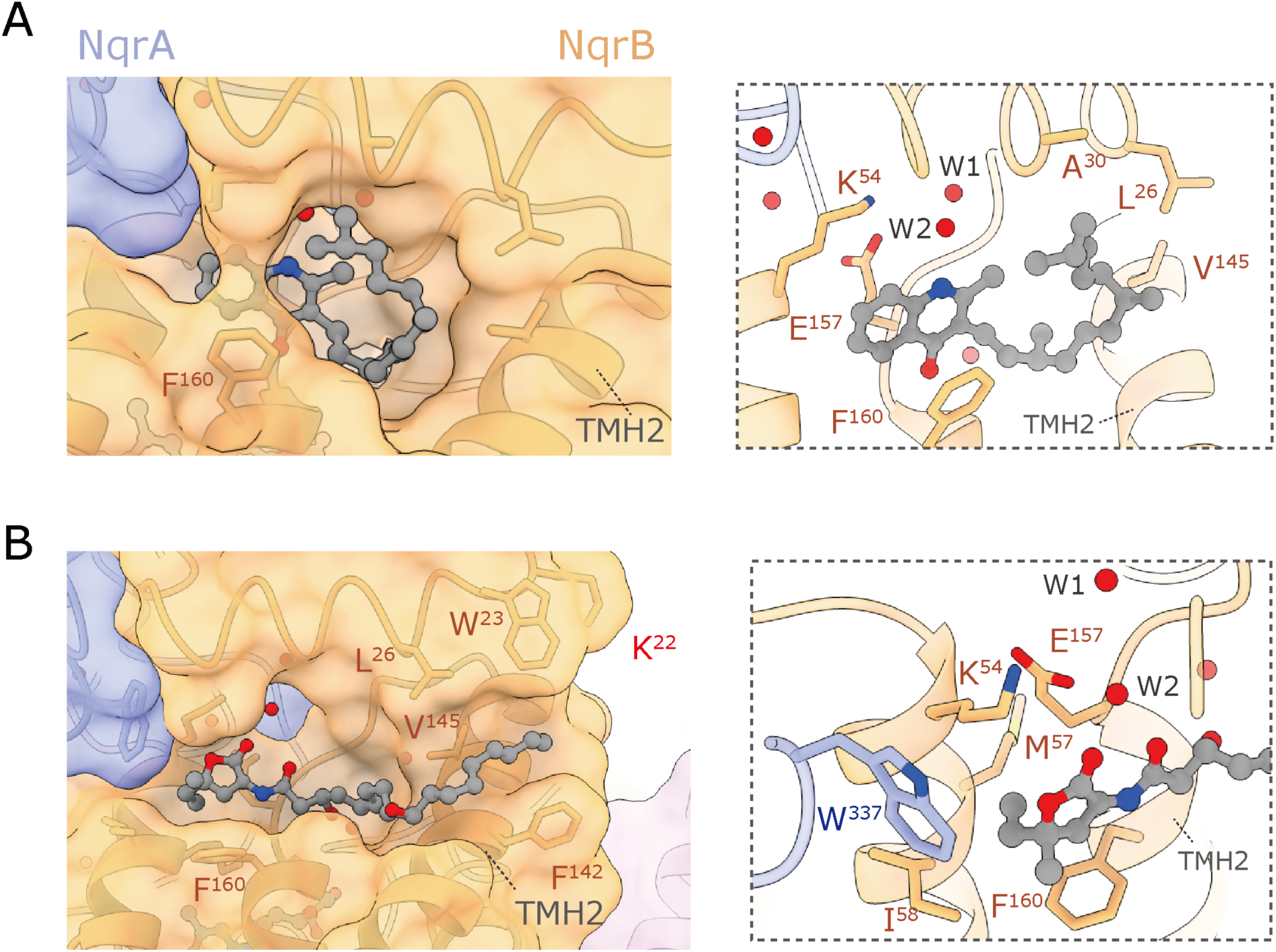
The binding manner of aurachin D-42 and korormicin A in NqrB. (**A**) Close-up view of the binding site of aurachin D-42. NqrB-Phe160 may be involved in a π-stacking interaction with the quinolone ring, which locks aurachin D-42 inside the binding cavity. (**B**) Close-up view of the binding site of korormicin A. The steric obstruction between the alkyl branches (5-CH_3_/C_2_H_5_) on the lactone ring and the cavity wall formed by NqrB-Met57 and -Ile58 on TMH 1, NqrB-Phe160 on the TMH 3, and NqrA-Try337 may fix the conformation of the lactone ring. The alkyl side chain extends toward a hydrophobic cleft composed of Trp23, Leu26, Phe142, and Val145 of NqrB.

**Extended Data Figure 8:**
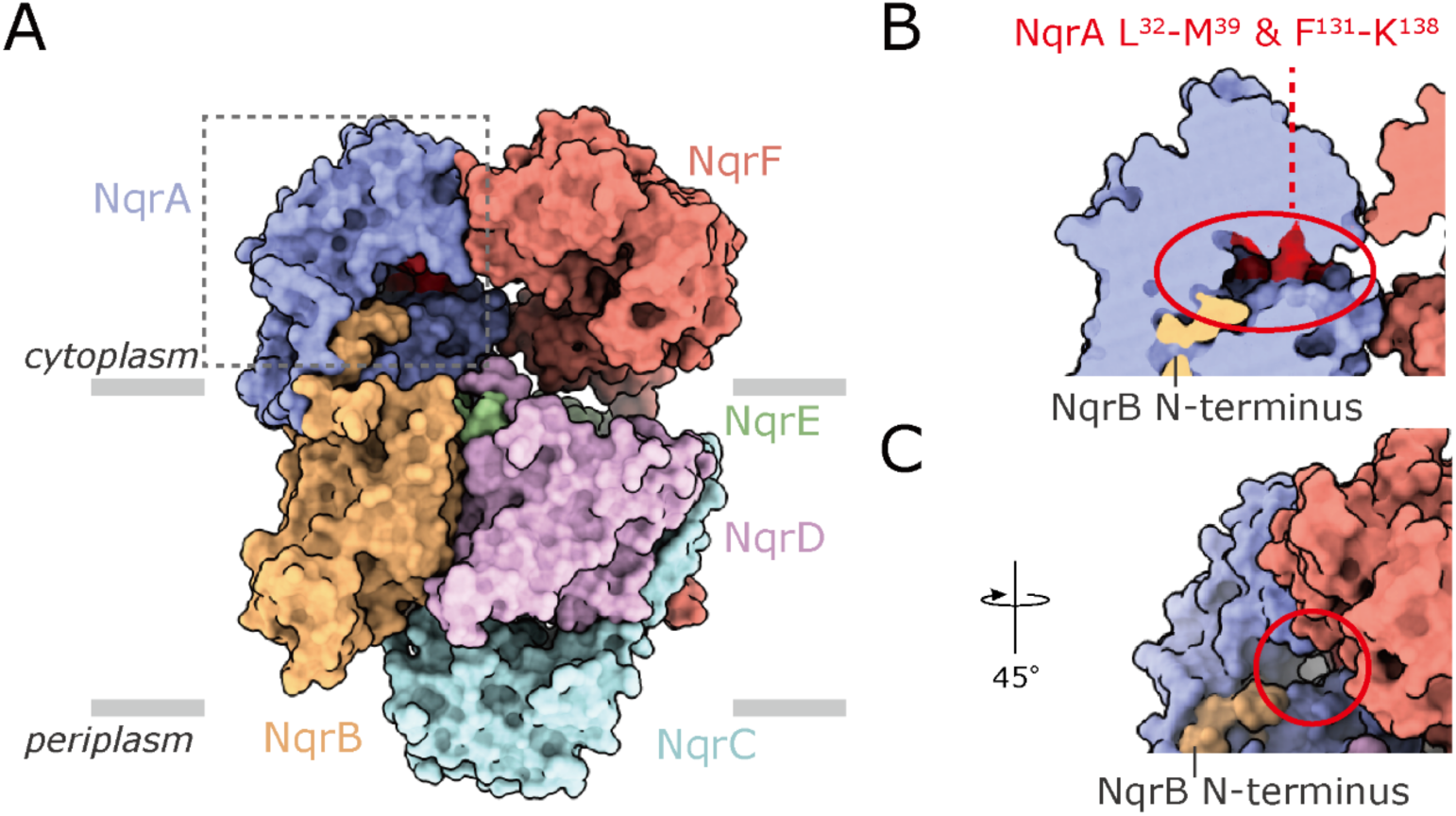
The binding site of the UQ head-ring in the cryo-EM structure. The site is indicated by a *red circle* in the current cryo-EM structure of Na^+^-NQR without bound inhibitor.

