## Supplementary Information for "Cryo-EM structures of Na^+^-pumping NADH-ubiquinone oxidoreductase from *Vibrio cholerae*"

**Supplementary Figure 1**

The binding sites of inhibitors and the UQ head-ring identified by photoaffinity labeling

**Supplementary Figure 2**

*N*-Acyl-*N*-alkyl sulfonamide chemistry

**Supplementary Figure 3**

Structure-inhibition relationship of korormicin A derivatives

**Supplementary Tables 1 and 2**

Details for the data processing and statistics for the maps and modeling

**Supplementary Table 3**

Comparison of the r.m.s.d values for each subunit between the cryo-EM and crystallographic structures

**Supplementary Discussion**

Equilibrium binding model of $^{125}$I-incorporated inhibitor and competitor

(A)

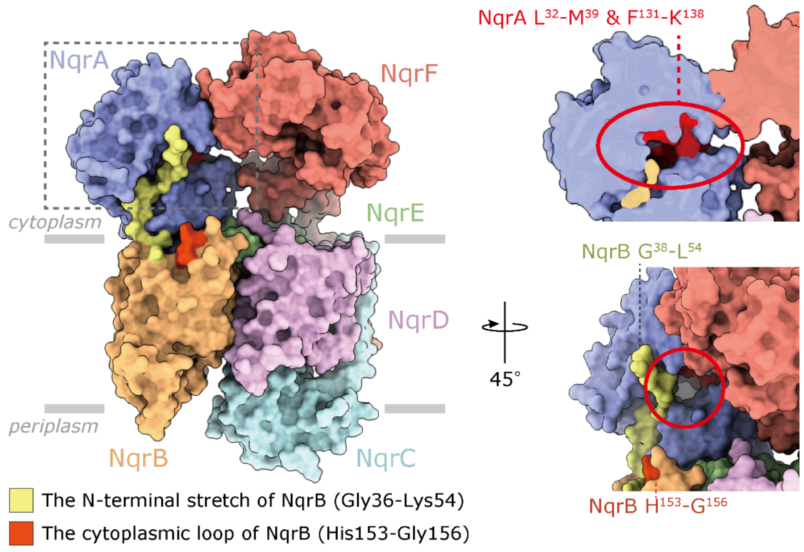


(B)

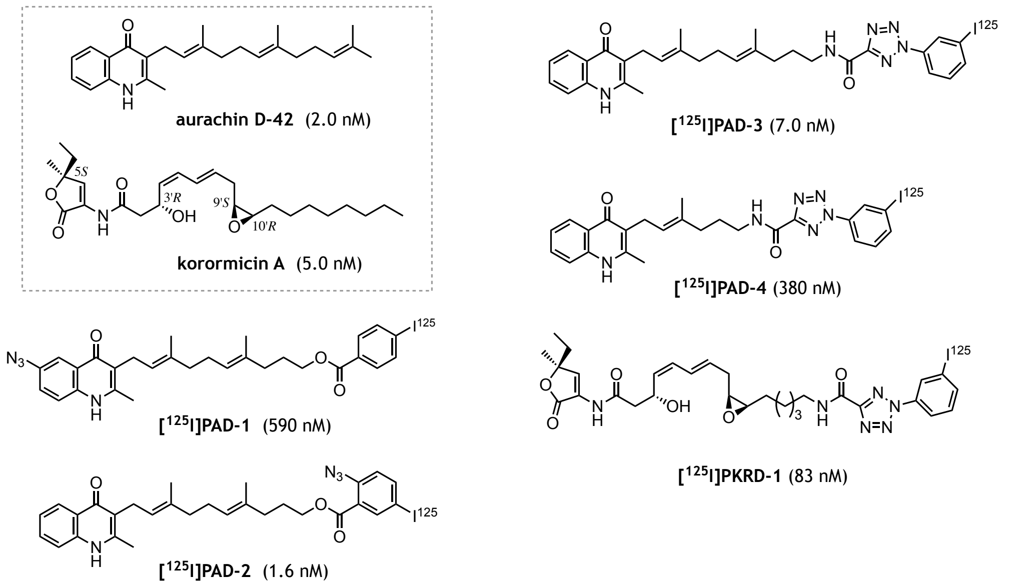


**Supplementary Figure 1**: **The binding sites of inhibitors and the UQ head-ring identified by photoaffinity labeling.** (**A**) The binding sites of inhibitors in NqrB and the UQ head-ring in NqrA are shown in the X-ray crystallographic structure (*16*, PDB ID: 4P6V). The korormicin A derivatives ([$^{125}$I]PKRD-1) and aurachin D derivatives ([$^{125}$I]PAD-1−[$^{125}$I]PAD-4) bind to the region (Try23−Lys54 in *yellow*) in the protruding N-terminal stretch starting with TMH 1 of NqrB and/or a part of the cytoplasmic loop (His153−Gly156 in *red*) connecting TMHs 2–3 of NqrB. The UQ head-ring binds to the cytoplasmic region of NqrA (Leu32−Met39 and Phe131−Lys138, indicated by a red circle). (**B**) The structures of the photoreactive inhibitors, which were used in the previous studies (*18* and *19*), are shown. The average $IC_{50}$ value of each inhibitor, which were determined with 1.0 nM $Na^+$-NQR, are shown in the parentheses.

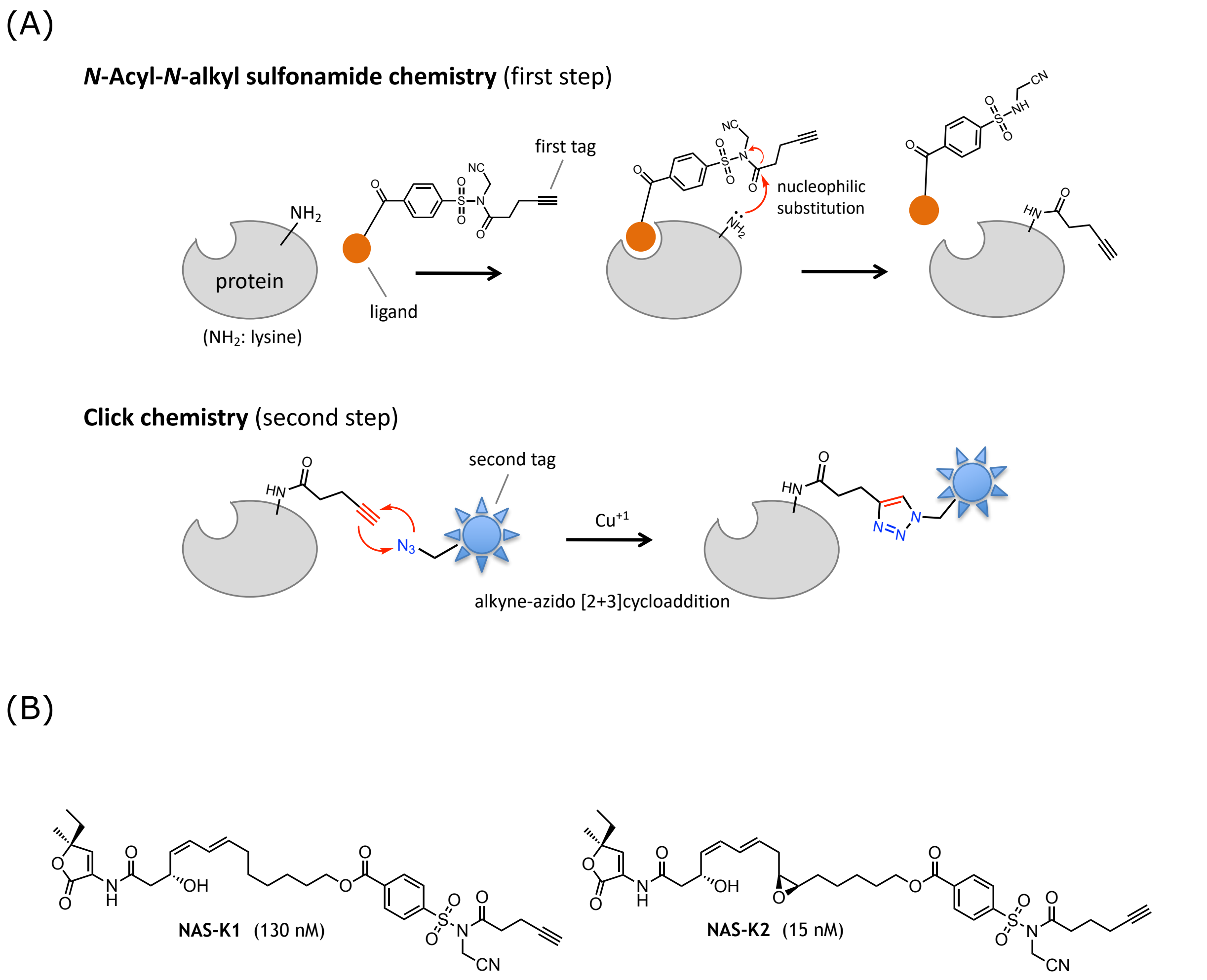


**Supplementary Figure 2: *N*-Acyl-*N*-alkyl sulfonamide chemistry.** (**A**) *N*-Acyl-*N*-alkyl sulfonamide chemistry is schematically shown. A first tag attached to the sulfonamide moiety is introduced to lysine via *N*-acyl-*N*-alkyl sulfonamide chemistry (*27*). Then, the reacted lysine can be identified by proteomic analyses after a second tag (e.g. fluorescent tag and biotin) is introduced to the modified lysine via click chemistry. (**B**) Structures of NAS-K1 and NAS-K2 used in the previous study (*21*).

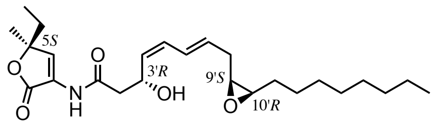


**korormicin A** (IC$_{50}$ = 5.0 nM)

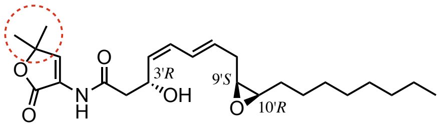


**compound D1** (IC$_{50}$ = 9.4 nM)

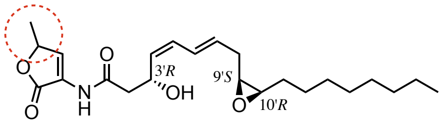


**compound D2** (IC$_{50}$ = 92 nM)

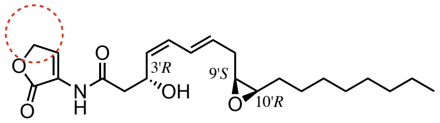


**compund D3** (IC$_{50}$ = 2700 nM)

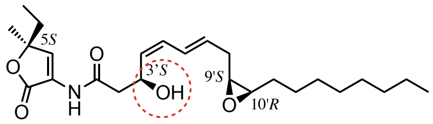


**compound D4** (IC$_{50}$ = 270 nM)

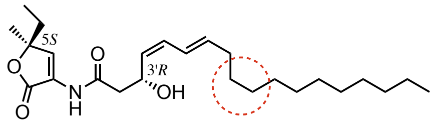


**compound D5** (IC$_{50}$ = 23 nM)

**Supplementary Figure 3: Structure-inhibition relationship of korormicin A derivatives.** Structure-inhibition relationship of korormicin A derivatives (*19*). The IC$_{50}$ value is the molar concentration needed to reduce the NADH-UQ$_1$ oxidoreductase activity of the isolated Na$^+$-NQR (1.0 nM) by 50%.

**Supplementary Table 1.** Statistics for Cryo-EM data, refinement, and validation of $Na^+$-NQR.

| | | $Na^+$-NQR | |
|---|---|---|---|
| **Data collection and processing** | | | |
| Magnification | | 81,000 | |
| Voltage (kV) | | 300 | |
| Total dose ($e^-/Å^2$) | | 65 | |
| Defocus range (μm) | | -0.8 to -2.0 | |
| Pixel size (Å) | | 0.88 | |
| Symmetry imposed | | C1 | |
| # of movies | | 9,027 | |
| # of Initial particles | | 2,570,565 | |
| **States** | **1** | **2** | **3** |
| EMDB ID | 33242 | 33243 | 33244 |
| PDB ID | 7XK3 | 7XK4 | 7XK5 |
| # of Final particle | 82,790 | 80,882 | 72,234 |
| Resolution (Å) FSC =0.143 | 3.1 | 3.1 | 3.1 |
| **Model statistics** | | | |
| Model resolution (Å) FSC =0.5 | 3.1 | 3.1 | |
| Model composition | | | |
| Non-hydrogen atoms | 14,847 | 14,835 | 14,835 |
| Residues | 1,894 | 1,894 | 1,894 |
| Ligands | FMN: 2, FES: 2, RBF: 1. FAD: 1, PEE: 1, CA: 1, LMT: 2, | FMN: 2, FES: 2, RBF: 1. FAD: 1, PEE: 2, CA: 1, LMT: 2, | FMN: 2, FES: 2, RBF: 1. FAD: 1, PEE: 1, CA: 1, LMT: 2, |
| Waters | 0 | 0 | 0 |
| Bond length (Å) | 0.006 | 0.006 | 0.005 |
| Bond angles (°) | 0.800 | 0.846 | 0.787 |
| Clash score | 6.16 | 5.77 | 5.60 |
| MolProbity score | 1.56 | 1.43 | 1.51 |
| EMRinger score | 3.79 | 2.68 | 2.78 |
| Rotamer outliers (%) | 0.35 | 0.47 | 0.45 |
| Ramachandran plot | | | |
| Favored (%) | 96.55 | 97.40 | 96.76 |
| Allowed (%) | 3.29 | 2.50 | 3.13 |
| Outlier (%) | 0.16 | 0.11 | 0.11 |

**Supplementary Table 2.** Statistics for Cryo-EM data, refinement, and validation of $Na^+$-NQR$^{AD42}$ and $Na^+$-NQR$^{KA}$.

| | $Na^+$-NQR$^{AD42}$ | $Na^+$-NQR$^{KA}$ |
|---|---|---|
| **Data collection and processing** | | |
| Magnification | 81,000 | |
| Voltage (kV) | 300 | |
| Total dose ($e^-$/Å$^2$) | 60 | |
| Defocus range (μm) | -0.8 to -2.0 | |
| Pixel size (Å) | 0.88 | |
| Symmetry imposed | C1 | |
| # of movies | 11,430 | 2,814 |
| # of Initial particles | 2,944,013 | 721,914 |
| EMDB ID | 33245 | 33246 |
| PDB ID | 7XK6 | 7XK7 |
| # of Final particle | 54,284 | 50,444 |
| Resolution (Å) FSC =0.143 | 3.0 | 2.9 |
| **Model statistics** | | |
| Model resolution (Å) FSC =0.5 | 3.0 | 3.0 |
| Model composition | | |
| Non-hydrogen atoms | 15,124 | 15,187 |
| Residues | 1,919 | 1,921 |
| Ligands | AUD:1, FMN: 2, FES: 2, RBF: 1. FAD: 1, PEE: 2, CA: 1, LMT: 2, | KRR: 1, FMN: 2, FES: 2, RBF: 1. FAD: 1, PEE: 2, CA: 1, LMT: 2, |
| Waters | 49 | 35 |
| Bond length (Å) | 0.006 | 0.012 |
| Bond angles (°) | 0.850 | 1.227 |
| Clash score | 4.69 | 4.96 |
| MolProbity score | 1.48 | 1.36 |
| EMRinger score | 3.68 | 4.09 |
| Rotamer outliers (%) | 0.58 | 0.57 |
| Ramachandran plot | | |
| Favored (%) | 96.37 | 97.48 |
| Allowed (%) | 3.42 | 2.36 |
| Outlier (%) | 0.21 | 0.05 |

**Supplementary Table 3**. The r.m.s.d values for each subunit between the cryo-EM-structure (this study) and crystallographic structure (*16*). The values were calculated using UCSF chimera. The distance between each Cα has been pruned by 5 Å.

| Subunit | r.m.s.d. (Å) |
|---|---|
| **NqrA** | 0.913 |
| **NqrB** | 1.349 |
| **NqrC** | 1.125 |
| **NqrD** | 2.107 |
| **NqrE** | 1.843 |
| **NqrF** | 1.317 |

**Supplementary Discussion:**

**Equilibrium binding model of $^{125}$I-incorporated inhibitor and competitor**

Based on the idea of two different conformations of a single binding cavity, we produced a new equilibrium binding model to explain competitive behavior between $^{125}$I-incorporated inhibitor (**[$^{125}$I]I**) and competitor (**C**), as shown in Supplementary Figure 4A.

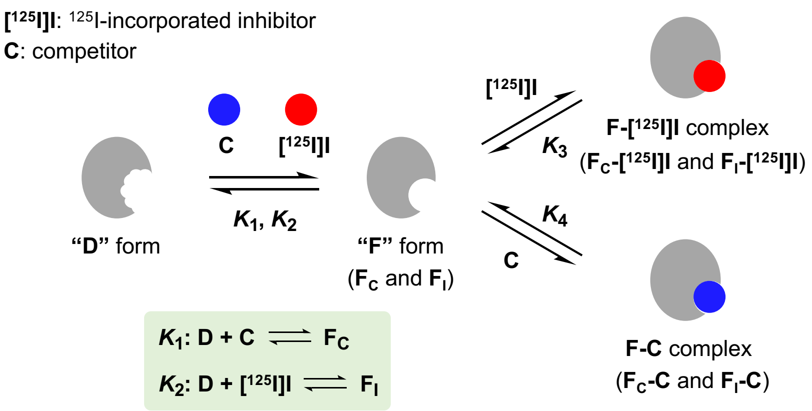


**Supplementary Figure 4A**. Schematic presentation of the equilibrium binding model based on two different conformations of a single binding cavity ("**D**" and "**F**" represent the disordered and fixed conformations, respectively).

Since the radioactivity incorporated into the enzyme by photolysis is proportional to the concentration of the enzyme-$^{125}$I-incorporated inhibitor complex (**F-[$^{125}$I]I** complex) regardless of the labeling reaction yields, we estimated the concentrations of **F-[$^{125}$I]I** complex by this equilibrium model. The "**D**" and "**F**" forms stand for the *disordered* and *fixed* conformations, respectively, of the binding cavity in the NqrB subunit. As the conformation of the cavity varies slightly depending on individual bound inhibitors, the **F** form takes two conformations in the strict sense: $\mathbf{F_C}$ (competitor-bound conformation) and $\mathbf{F_I}$ ([$^{125}$I]I-bound conformation). Therefore, regarding the equilibrium constants $K_1$ and $K_2$, we consider the following three cases (models 1–3). In the model 1, not only the conformations of $\mathbf{F_I}$ and $\mathbf{F_C}$ but also $K_1$ and $K_2$ are identical. This case corresponds to the competition between $^{125}$I-incorporated inhibitor and its non-radioactive (cold) analog (*18*). In the model 2, the conformations of $\mathbf{F_I}$ and $\mathbf{F_C}$ are identical but $K_1$ and $K_2$ are different. In the model 3, the conformations of $\mathbf{F_I}$ and $\mathbf{F_C}$ and the equilibrium constants $K_1$ and $K_2$ are both different. This case corresponds to the competition between $^{125}$I-incorporated inhibitor and a different type of inhibitor. Since the models 2 and 3 are extended cases of the model 1, we first numerically solved the equilibrium equations based on the model 1 ($K_1 = K_2$). The equilibrium compositions were calculated based on the kinetic simulations for a sufficiently long time by COMSOL Multiphysics® (COMSOL) under varying conditions. In the initial conditions of the simulation, the system contains only **D**, **[$^{125}$I]I**, and **C**.

For the current simulation, we set $K_1$, $K_2$, $K_3$, and $K_4$ to 100,000, 100,000, 1,000,000, and 1,000,000 $mM^{-1}$ and the total concentration of **[$^{125}$I]I** to 10 nM under different total concentrations of both the enzyme and competitor (**C**). Note that the total concentrations of **[$^{125}$I]I** (10 nM) and the enzyme (0.90, 9.0, 90, and 900 nM) were set to those used in the previous photoaffinity labeling experiments (*18*) throughout the following simulations. Based on these assumptions, the changes of the concentration of individual species (**D**; enzyme in D-form, **F**; enzyme in F-form, **F-[$^{125}$I]I**; enzyme-$^{125}$I-incorporated inhibitor complex, **F-C**; enzyme-competitor complex) against the initial concentration of competitor ([**C**]$_0$) for four different enzyme concentrations were simulated, as shown in Supplementary Figure 4B.

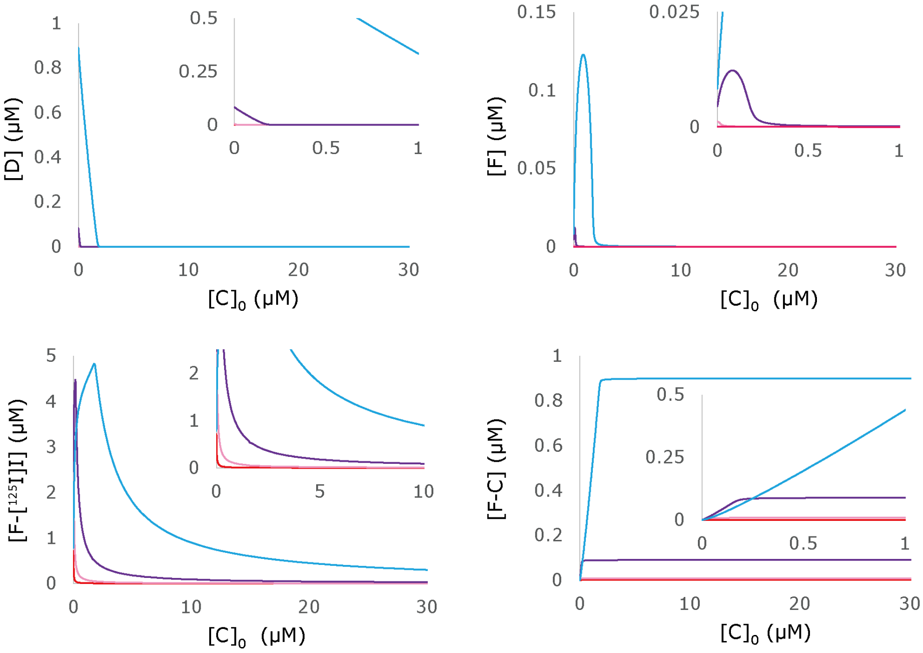


**Supplementary Figure 4B**. Changes of the concentration of individual species (enzyme in D-form (**D**), enzyme in F-form (**F**), enzyme-$^{125}$I-incorporated inhibitor complex (**F-[$^{125}$I]I**), enzyme-competitor complex (**F-C**)) against the initial concentration of competitor ([**C**]$_0$) with different enzyme concentrations. The concentration of $Na^+$-NQR: 0.9 (*red*), 9.0 (*pink*), 90 (*purple*), and 900 nM (*blue*).

Then, the concentrations of **F-[$^{125}$I]I** complex as a function of the concentrations of added competitor (**C**) were normalized by the concentration of **F-[$^{125}$I]I** when [C]$_0$ = 0, so that the value at zero added competitor is one (Supplementary Figure 4C); values below one indicate competitive suppression of the formation of **F-[$^{125}$I]I** complex, while values above one indicate enhancement. Our model can account for the consecutive changes of the effects of the competitor, from enhancement to suppression, as the concentration of the competitor increases in the case of a high concentration of the enzyme (900 nM). The enhancing effect became less clear with decrease in the enzyme concentrations. These simulation results are consistent with the experimental ones (*18*).

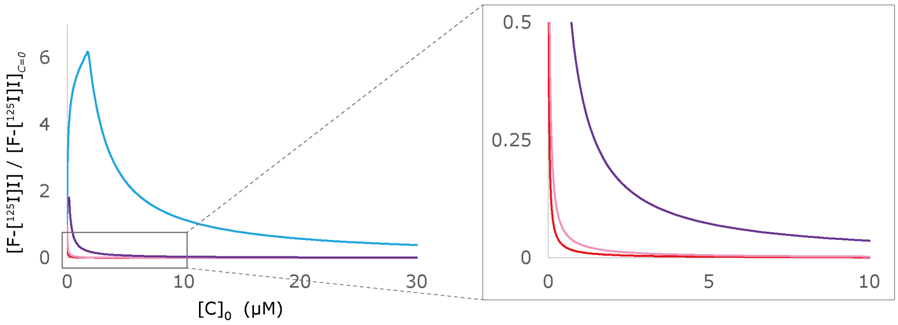


**Supplementary Figure 4C**. The concentration of **F-[$^{125}$I]I** complex as a function of the initial concentration of competitor (**C**), normalized so that the value at zero added competitor is one. The concentration of $Na^+$-NQR: 0.9 (*red*), 9.0 (*pink*), 90 (*purple*), and 900 nM (*blue*).

For reference, changes of the concentrations of **F-[$^{125}$I]I** complex when the parameters ($K_1$–$K_4$) are varied are shown in Supplementary Figure 4D. Similar tendencies in the changes of **F-[$^{125}$I]I** complex with those shown in Supplementary Figure 4C were observed.

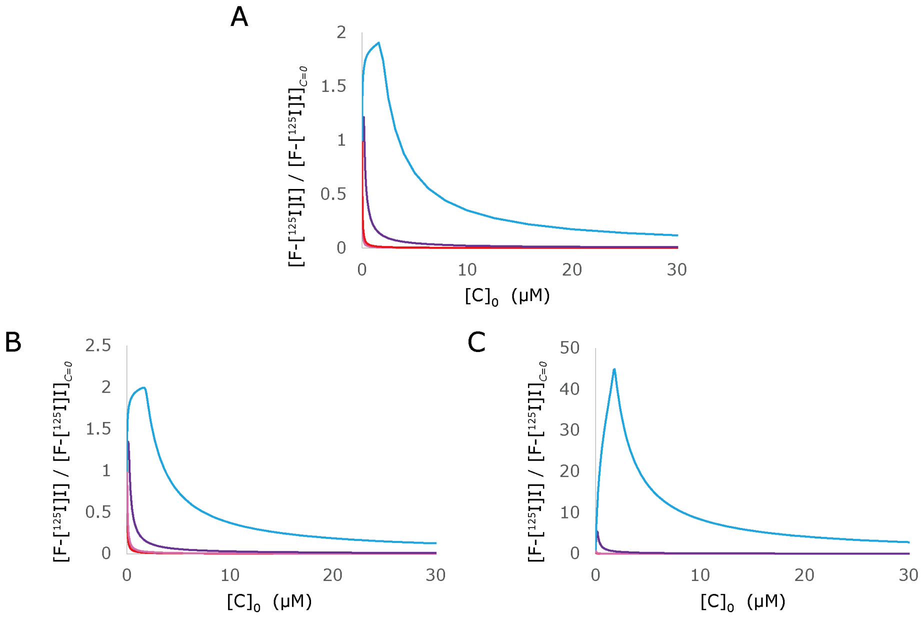


**Supplementary Figure 4D**. The concentration of **F-[$^{125}$I]I** complex as a function of the initial concentration of competitor (**C**). The simulations were conducted under three different conditions. Panel A: $K_1$, $K_2$, $K_3$ and $K_4$ were set to 100,000, 100,000, 10,000,000, and 10,000,000 $mM^{-1}$, respectively. Panel B: $K_1$, $K_2$, $K_3$ and $K_4$ were set to 10,000, 10,000, 1,000,000, and 1,000,000 $mM^{-1}$, respectively. Panel C: $K_1$, $K_2$, $K_3$ and $K_4$ were set to 1,000,000, 1,000,000, 1,000,000, and 1,000,000 $mM^{-1}$, respectively. The concentration of $Na^+$-NQR: 0.9 (*red*), 9.0 (*pink*), 90 (*purple*), and 900 nM (*blue*).

Next, we conducted the simulations according to model 2 ($K_1 \neq K_2$, but $\mathbf{F_C}$ is identical to $\mathbf{F_I}$). Based on the results of model 1, the $K_1$, $K_2$, $K_3$, and $K_4$ were set to 100,000, 100,000, 1,000,000, and 1,000,000 $mM^{-1}$, respectively. The consecutive changes of the effects of the competitor, from enhancement to suppression, were observed (Supplementary Figure 4E), as seen in the simulations based on model 1 (Supplementary Figure 4C).

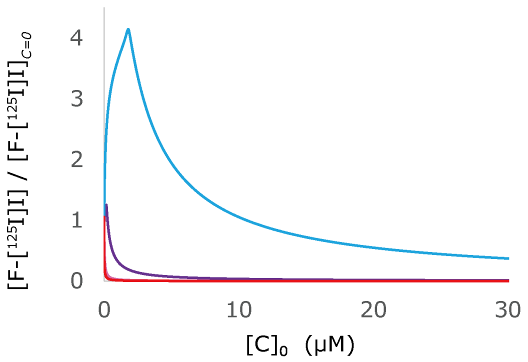


**Supplementary Figure 4E.** The concentration of **F-[[125]I]I** complex as a function of the initial concentration of competitor (**C**), calculated based on the model 2. The $K_1$, $K_2$, $K_3$ and $K_4$ were set to 100,000, 200,000, 1,000,000, and 1,000,000 mM$^{-1}$, respectively. The concentration of $Na^+$-NQR: 0.9 (*red*), 9.0 (*pink*), 90 (*purple*), and 900 nM (*blue*).

Finally, we conducted simulations according to model 3 ($K_1 \neq K_2$, and $\mathbf{F_C}$ is not identical to $\mathbf{F_I}$). As the conformations of the $\mathbf{F_C}$ and $\mathbf{F_I}$ forms are different in this case, $K_3$ and $K_4$ increase twofold as follows:

$$K_{3\text{-}1}\text{: } \mathrm{F_C} + [^{125}\mathrm{I}]\mathrm{I} \rightleftharpoons \mathrm{F_C} - [^{125}\mathrm{I}]\mathrm{I}$$

$$K_{3\text{-}2}\text{: } \mathrm{F_I} + [^{125}\mathrm{I}]\mathrm{I} \rightleftharpoons \mathrm{F_I} - [^{125}\mathrm{I}]\mathrm{I}$$

$$K_{4\text{-}1}\text{: } \mathrm{F_C} + \mathrm{C} \rightleftharpoons \mathrm{F_C} - \mathrm{C}$$

$$K_{4\text{-}2}\text{: } \mathrm{F_I} + \mathrm{C} \rightleftharpoons \mathrm{F_I} - \mathrm{C}$$

We conducted the simulation by setting $K_1$, $K_2$, $K_{3\text{-}1}$, $K_{3\text{-}2}$, $K_{4\text{-}1}$, and $K_{4\text{-}2}$ to 100,000, 200,000, 1,000,000, 2,000,000, 1,000,000, and 2,000,000 mM$^{-1}$, respectively. Again, similar tendencies in the changes of the concentration of **F-[[125]I]I** complex (a sum of **$\mathbf{F_C}$-[[125]I]I** and **$\mathbf{F_I}$-[[125]I]I**) with those simulated by model 1 were observed (Supplementary Figure 4F).

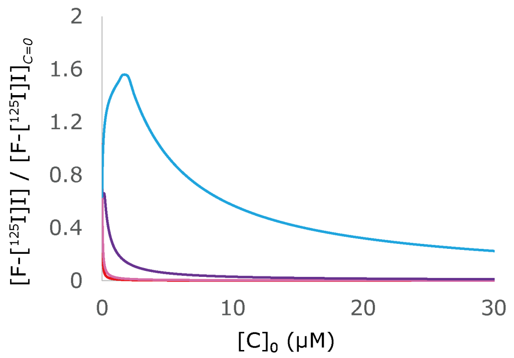


**Supplementary Figure 4F**. The concentration of **F-[[125]I]I** complex as a function of the initial concentration of competitor (**C**), calculated based on the model 3. The $K_1$, $K_2$, $K_{3\text{-}1}$, $K_{3\text{-}2}$, $K_{4\text{-}1}$, and $K_{4\text{-}2}$ were set to 100,000, 200,000, 1,000,000, 2,000,000, 1,000,000, and 2,000,000 mM$^{-1}$, respectively. The concentration of $Na^+$-NQR: 0.9 (*red*), 9.0 (*pink*), 90 (*purple*), and 900 nM (*blue*).

Altogether, the unusual competitive behavior observed in the previous photoaffinity labeling study (*18*) can be accounted for by the equilibrium model based on the idea of two different conformations of a single binding cavity. The extents of enhancing or suppressing effects vary depending on the parameters ($K_1$–$K_4$) that are determined by the individual chemical nature of $^{125}$I-incorporated inhibitors and competitors used. Therefore, small differences in the effects among different pairs of $^{125}$I-incorporated inhibitor and competitor, which were observed in the labeling experiments (*18*), would easily be accounted for.
